# From deployment to functioning ecosystems: assessing taxonomic richness and community composition on modular shellfish reefs using environmental DNA metabarcoding

**DOI:** 10.64898/2026.10.26.762076

**Authors:** Valerio Tettamanti, Timothy L. Staples, Benjamin Mos, Robbie Porter, Ben L. Gilby, Karen L. Cheney

## Abstract

Ecological restoration success is commonly assessed against historical reference states that are often altered, poorly defined, or absent, particularly in marine systems. Reliance on taxon richness alone can further obscure changes in community structure, successional trajectories, and the emergence of functionally important taxa at restoration sites. We assessed biodiversity recovery on modular shellfish reef structures using environmental DNA (eDNA) metabarcoding, combining taxon-specific richness models with multivariate community composition analyses across a four-year age gradient (9, 21, and 42 months). Genus richness increased rapidly following reef deployment but differed strongly among taxon groups. Dominant groups, particularly Mollusca, showed similar richness in nine-month-old and 42-month-old reefs; crustaceans and fishes also showed high, consistent richness early in development. Five of nine taxon groups displayed significant richness changes through time, with increasing (flatworms and macroalgae) and decreasing (cnidarians, sponges, and roundworms) trajectories. Community composition showed pronounced age-related structuring: younger reefs were compositionally distinct from older reefs, and within-age variability declined markedly with age, consistent with convergent successional assembly. Trait-based patterns indicated shifts towards non-benthic crustaceans, larger-bodied fish, and free-living flatworms in older reefs. These results show that modular shellfish structures are rapidly colonised by a wide taxonomic range within 9 months, producing early richness gains. However, the identity and relative representation of those taxa continue to change, generating structured compositional shifts undetectable from richness alone. Integrated richness–composition frameworks focused on ecological trajectories provide a more robust basis for evaluating restoration success than strict reference states. Environmental DNA metabarcoding offers a scalable, non-invasive monitoring approach capable of capturing both early richness gains and longer-term community assembly dynamics in data-limited marine systems.

## Introduction

The goal of ecological restoration is to begin or support the restoration of ecosystems that have been harmed, degraded, or destroyed, with the goal of fully restoring their structure, functions, and biological diversity (Balensiefer et al., 2004). To do so, projects have commonly aimed to return a system towards a historic or reference state (Aronson and Alexander, 2013; Benayas et al., 2009; Hallett et al., 2013; Hobbs and Harris, 2001; Suding, 2011; Suding et al., 2004). However, complete restoration requires not only recovery of basic ecological structure, but the re-establishment of fully functioning assemblages of native taxa and the ecological processes they support (Suding, 2011). Although reference conditions can provide useful benchmarks for setting restoration goals, ecosystems are dynamic and historically contingent, particularly in environments subject to long-term exploitation and environmental change (Myers and Worm, 2003; Pauly, 1995; Pinnegar and Engelhard, 2008). As a result, restoring a single historical reference state is increasingly unattainable (Duarte et al., 2009; Pinnegar and Engelhard, 2008).

In the absence of a known or achievable reference state, restoration success can be assessed through approaches that integrate ecological theory with explicitly articulated values of biodiversity, or alternative ecosystem states (Brewer and Menzel, 2009; Hagger et al., 2025; Suding et al., 2004). Such frameworks define success in terms of desired contemporary ecological and social outcomes, including the recovery of ecosystems functions, biodiversity gains and cultural connections (Duarte et al., 2009; Smith et al., 2022). Importantly, these approaches provide a transparent basis for prioritising functional and biodiversity outcomes where historical benchmarks are uncertain, inappropriate, or unattainable. A key remaining challenge is therefore defining what constitutes value and how best to measure it, whether through biodiversity metrics, functional recovery, or ecosystem service provision.

Shellfish reefs - particularly those formed by oysters - once dominated coastal regions, estuaries, and embayments worldwide. An estimated 85% of global oyster reefs have been lost through the combined effects of overfishing, lime quarrying, sedimentation, disease, and coastal development (Beck et al., 2011; Lotze et al., 2006; Ogburn et al., 2007). This widespread decline has resulted in extensive habitat degradation, loss of fisheries income, and erosion of cultural services (Beck et al., 2011; zu Ermgassen et al., 2013). For First Nations Peoples, shellfish have long held and continue to hold cultural and economic significance, provided food security, and underpinned environmental sustainability (Gibbs et al., 2024). Consequently, the restoration of oyster reefs represents both an ecological priority and a culturally meaningful pathway for recovering lost biodiversity and ecosystem services (Gibbs et al., 2023; Grabowski and Peterson, 2007).

Against this backdrop of widespread ecological loss and enduring cultural significance, oyster reef restoration provides a valuable case study for examining how restoration success and ecological value can be defined and measured and offers a compelling system in which to investigate how ecological communities reassemble through successional processes following intervention. Shellfish reef restoration is often framed in the context of succession and the “field of dreams” restoration hypothesis, whereby reinstating habitat structure can facilitate natural community assembly processes (Hilderbrand et al., 2005; Palmer et al., 1997). In practice, restoration typically involves rebuilding habitat structure such as oyster shells or rock to restore benthic complexity and foundational processes. While such habitat provide settlement surfaces for sessile reef-forming taxa they also create structure and resource availability for higher trophic and mobile community members, with subsequent community assembly driven largely by natural colonisation and other broader ecological processes.

The underlying assumption is that the most ecologically valuable and functionally effective community will emerge, following deterministic theories of succession. In reality, both deterministic and stochastic processes interact to form communities over multiple temporal scales (Fukami and Nakajima, 2011; Perring et al., 2015). However, observing restoration sites whose community compositions converge over time can indicate that deterministic processes are dominant, and that some kind of functional attractor state exists (Walker et al., 2010). This chronosequential approach to restoration monitoring is popular in terrestrial systems but is only useful where restoration is rapid. Oyster reefs can regain some service provision rapidly (i.e. shoreline protection) while other services, such as fishery benefits, can take longer (Baggett et al., 2015; Smith et al., 2022).

If restoration success is defined in terms of ecological trajectories and functional recovery, robust monitoring becomes essential for assessing how communities reorganise and regain functionality following establishment. Long-term monitoring will also detect deviations or bottlenecks in recovery that may require adaptive intervention, and may also improve the efficacy of restoration intervention, ensuring efficient use of financial and ecological investment. However, long-term monitoring of restored ecosystems remains a significant challenge, and there is not yet a widely adopted global set of standardised metrics applied across consistently across restoration systems (Ruiz-Jaen and Mitchell Aide, 2005). This challenge is particularly acute in marine environments, where restoration success is commonly assessed using habitat-forming foundational species, and comprehensive, whole community biodiversity monitoring is rarely implemented, especially for cryptic or small-bodied invertebrates that may evade conventional surveys (Eger et al., 2022; Saenz-Agudelo et al., 2022).

Environmental DNA (eDNA) metabarcoding has emerged as a powerful, non-invasive method for generating a shallow-but-wide census of entire communities. By detecting DNA shed by organisms in their environment, eDNA enables rapid and non-destructive sampling of taxa from soil microbes to mammals. When interpretated in a functional context, such community-wide detection can provide insights into the recovery of key taxa and processes underpinning ecosystem function (Beng and Corlett, 2020; Gold et al., 2021). However, eDNA data typically only provides presence-only data, with limited insights into abundance or population structure. In aquatic environments, signals can vary spatially and temporally due to water movement, degradation rates and shedding dynamics. In addition, uneven bioinformatic pipelines and incomplete reference databases lead to potential misidentification or gaps in detection. Nevertheless, because eDNA can generate rapid, community-wide biodiversity assessments with higher detection sensitivity compared to traditional taxonomic surveys it represents a feasible and ecologically informative approach for tracking community trajectories on restored shellfish reefs, providing a compelling rationale for evaluating its performance in this context.

Here, we assessed the use of eDNA metabarcoding as a tool for capturing a broad, community-wide census of biodiversity on modular shellfish restoration structures. Specifically, we examined changes in community composition on modular Robust Oyster Baskets (ROBs) that had been deployed in the field for 9, 21 and 42 months. This approach enabled the detection of a wide spectrum of taxa across multiple kingdoms and allowed us to test whether post-restoration communities converged through deterministic successional processes. By tracking temporal changes in community composition, we aimed to determine timescales over which restored reefs may begin to provide measurable biodiversity benefits and characterise successional patterns.

## Methods

### eDNA sampling from restoration structures

We used Robust Oyster Baskets (ROBs) restoration structures designed and constructed by the not-for-profit organisation OzFish Unlimited Ltd (NSW, AUS). These structures are triangular prism-shaped steel-mesh containers (400 L × 400 W × 300 mm H; ‘ROB 400s’) filled with sterilised, recycled, locally sourced oyster shells (Fig. 1). ROBs were initially deployed in November 2020, November 2022, and November 2023 in linear arrays (lines 3–5 m apart; ROBs positioned approx. 0.3 m apart, Fig. 1), at an intertidal oyster lease off the coast of Myora Springs (−27.45684, 153.39528), North Stradbroke Island/Minjerribah, within Moreton Bay/Quandamooka, Australia. Over time, deployed ROBs became colonised by ambient marine communities, including rock oysters and other filter-feeding bivalves (Fig. 1).

**Figure 1.**
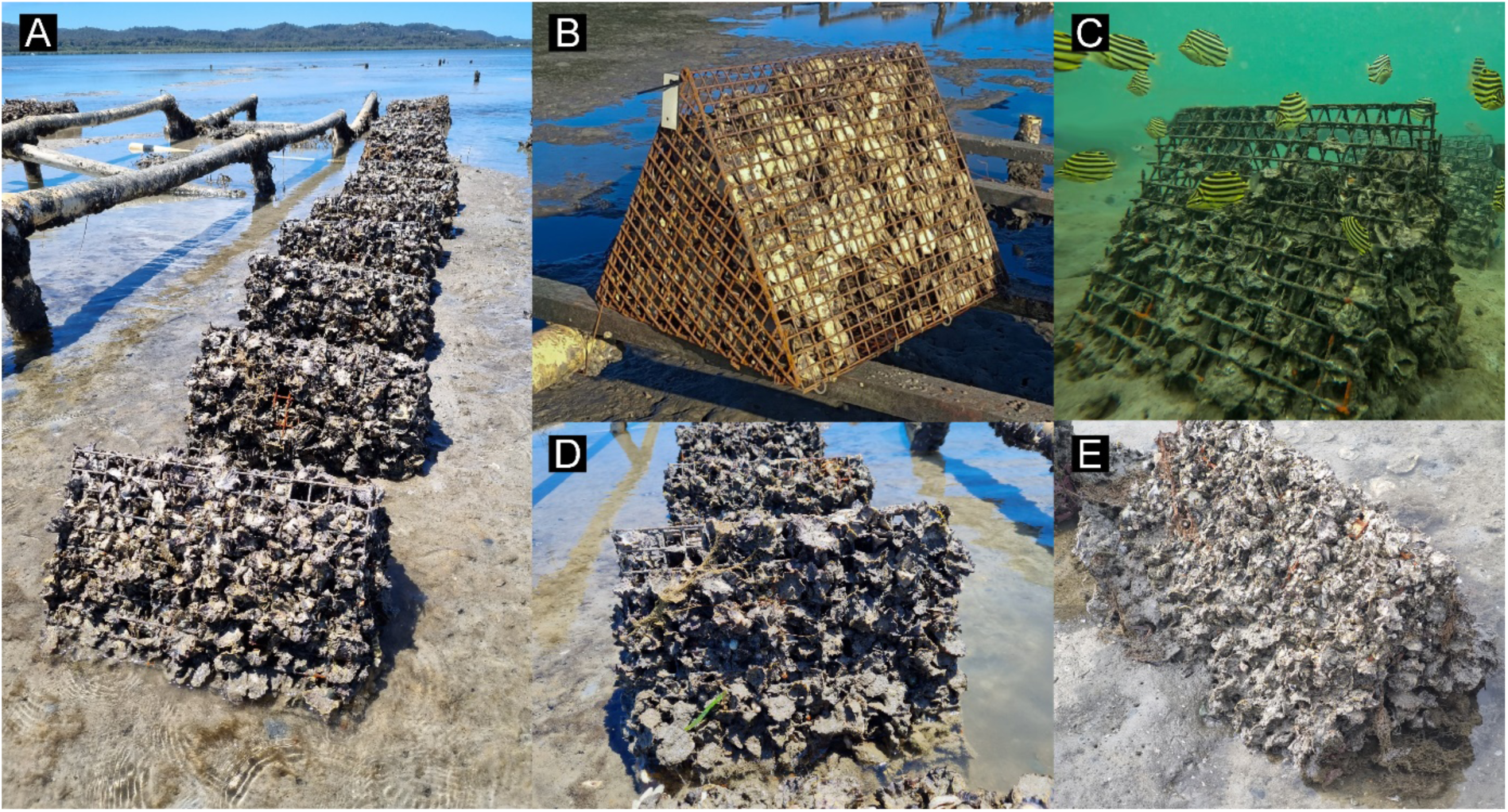
**A–E** Representative photographs of modular shellfish reef restoration structures, Robust Oyster Baskets (ROB). **A.** ROBs were deployed in lines in the intertidal zone in Moreton Bay/Quandamooka, Australia for 9–42 months. **B.** ROB prior to deployment, measuring 400 × 400 × 300 mm (L × W × H). Each ROB comprises a triangular prism-shaped steel-mesh cage filled with sterilised, recycled oyster shells, designed and constructed by not-for-profit OzFish Unlimited Ltd. **C.** ROB after ∼9 months deployment showing initial colonisation by shellfish and use as habitat by fish (high tide). **D.** ROB after ∼21 months deployment, showing substantial colonisation and growth by shellfish (low tide) and the beginnings of degradation of wire mesh due to corrosion. **E.** ROB after ∼36 months deployment, showing almost total colonisation by shellfish community (low tide), and showing similar appearance to the 42-months-deployed ROBs used in this study. Image attributions: R. Porter.

ROBs were retrieved from the field by hand after they had been deployed for different durations (∼9 months, n = 3; ∼21 months, n = 3; ∼42 months, n = 3). Due to logistical constraints, all ROBs could not be retrieved during the same period. The 42-month ROBs were retrieved on the 24/05/2024, while the 9 and 21 months ROBs were retrieved on the 29/08/2024. At low tide, ROBs were lifted off the lease by hand by researchers wearing gloves and immediately placed into round, lidded 110-L polyethylene tubs (internal dimensions: 400 mm H, 590 mm Ø). All tubs and lids were decontaminated at least one week prior to retrieval with 4% bleach solution, thoroughly rinsed, and kept sealed until use to minimize the risk of exogeneous DNA contamination. Once placed into the tubs, the ROBs were then transported to The University of Queensland’s Moreton Bay Research Station (MBRS) via a 30-minute boat trip for further processing.

On the day of collection, artificial saltwater was prepared using reverse osmosis (RO)-purified water and aquarium-grade marine salt (Tropic Marin, Wartenberg, Germany) in sterilised containers. On return to MBRS, artificial seawater was added to each previously described tub to fully submerge each ROB. Sterilized pumps, air tubes and air stones were used to circulate the water for a minimum of 1 h, while lids remained closed, allowing eDNA to be released into the water from the ROBs, while minimising the introduction of airborne or handling-derived contaminates. To monitor and account for potential contamination due to field handling and laboratory processing, a procedural negative control was included on both collection dates. An empty, sterilised tub was handled by researchers, transported to the field site, and processed identically to experimental samples, but without the inclusion of a ROB being added to the tub. DNA detected in the procedural negative control was used to identify and remove potential contaminants from the experimental samples prior to downstream analyses (see below).

Environmental DNA (eDNA) was collected from each ROB replicate using eDNA mini-kits from Wilderlab NZ Ltd (Wellington, New Zealand; https://www.wilderlab.co/), which included a sterile syringe, and a 30 mm × 1.2 μm cellulose acetate syringe filter with luer-lock inlet and outlet fittings. The kit also included sterile nitrile gloves, a 60 cc luer-lock syringe for drawing and pushing water through the filter membrane, and 350 μL of DNA/RNA Shield preservative (Zymo Research Cat. No. R1200-125) pre-filled into a 3 cc luer-lock syringe. We used 6 kits per ROB and 3 kits for each control, and aimed to filter 1 L of water per kit (total of 6 L per ROB and 3 L per control). The high number of sediments and particles in the water clogged the filter at lower volumes in some replicates (see Supplementary Tables S1 and S2). Following preservation, syringe filters were stored at room temperature and shipped to Wilderlab for downstream processing within two weeks of collection.

### DNA Extraction and Library Preparation

All laboratory processing was carried out by Wilderlab under their standard ECOSCAN protocol (v2.2.5; Wilkinson, 2025), summarised here. Lysate was recovered from each encapsulated filter with a sterile 3 mL syringe, held at −20 °C, and a 200 µL aliquot extracted on a Genolution Nextractor NX-48S using GD141 cartridges under default settings, with every extraction plate carrying at least one 200 µL DNase-free water blank (IDT) as an extraction control. Extraction quality assessment, adapter fusion, indexing and amplification were combined into single-step PCRs on an Applied Biosystems ProFlex, run in duplicate for each sample and blank. The protocol targeted a broad taxonomic range using fusion-tag assays spanning mitochondrial (12S, 16S), nuclear (18S, ITS) and chloroplast (rbcL, trnL) loci (Table S3), with each primer carrying Illumina P5/P7 adapters, the TruSeq forward sequencing primer site, a unique 8–9 bp index (differing from all others by ≥3 bp), and the locus-specific sequence. Each reaction contained 3 µL MyTaq 2× Red Mix (Bioline) with BSA at 2 mg mL⁻¹ (Sigma Aldrich), 0.5 µL of each primer (10 µM) and 1.5 µL template, cycled at 95 °C for 3 min followed by 38 cycles of 95 °C for 5 s, an assay-specific annealing step (Table S3) for 10 s, and 72 °C extension of 10 s. Reagent concentrations and cycle number (38) were identical across all assays.. Amplicons, blanks included, were then pooled, purified and size-selected with AMPure XP beads, using 0.9× and 1.2× bead ratios to bound the retained fragment size. Pool concentration was quantified on a Qubit 4 fluorometer (ThermoFisher) and normalised to 650 pM, and the library sequenced on an Illumina NextSeq 1000 using a NextSeq 1000/2000 P1 XLEAP-SBS cartridge spiked with 15% PhiX, run in a single direction for 200 cycles.

### ASV generation and taxonomic assignment

Sequence processing followed Wilderlab’s in-house pipeline (Wilkinson, 2025; Wilkinson et al., 2024). Reads were de-multiplexed in R (R Core Team, 2021) with insect (v 1.4.0; Wilkinson et al., 2018), trimmed, and resolved into exact amplicon sequence variants (ASVs) using DADA2 (Callahan et al., 2016). Taxonomy was assigned against a curated global reference built mainly from GenBank (Benson et al., 2013), BOLD (Ratnasingham and Hebert, 2007), and, for the UM assay only, the RDP database (v18; Cole et al., 2014). Any ASV matching a reference at 100% identity and coverage was resolved to its lowest common ancestor, i.e. to genus where more than one species matched and to family where more than one genus matched, while the low-resolution markers (BE, BU, BX, UM) were capped at genus regardless of match. ASVs longer than 50 bp with no exact match were then classified with SINTAX (Edgar, 2016) under a strict confidence cutoff of >0.99, again to a maximum of genus.

### Taxonomic harmonization and dataset filtering

We filtered the raw bioinformatic data to better reflect diversity of multicellular taxa of marine restoration importance. Dataset size at each step is shown at Table S4. First, we removed “root” entries that were unidentified DNA sequences. We then harmonized Genbank and RDP database entries using the World Register of Marine Species using the worrms package (Chamberlain and Vanhoorne, 2023). We retrieved WoRMS records for NCBI taxon IDs, returning taxonomic data for 2,597 NCBI IDs (70.2%). IDs without WoRMS records were largely bacterial groups and kingdom-level IDs for Eukaryota. We used the WoRMS harmonization to exclude taxa that were either not listed in WoRMS or not registered as occurring in either marine or brackish environments.

Given our focus on multicellular diversity and interest in taxa of importance for cultural, economic, and ecosystem services, we restricted analyses to taxa from the following groups: molluscs (Mollusca), crustaceans (Crustacea), cnidarians (Cnidaria; including jellyfish and coral), fish (Chordata; Actinopterygii and Chondrichthyes), flatworms (Platyhelminthes), roundworms (Nematoda), segmented worms (Annelida), sponges (Porifera) and macroalgae (Chlorophyta, Phaeophyta and Rhodophyta). Within these groups, we tested for potential contaminant taxa using control samples (Table S5). Contaminant taxa were defined as those present in at least one sample of both control rounds. We identified two contaminant taxa that were removed (Table S5). *Saccostrea* was flagged as a contaminant but was retained as it is a known member of oyster reefs. Finally, we removed control data from the dataset, retaining only eDNA samples from oyster ROBs. The harmonization and dataset filtering were conducted in R (R Core Team, 2021).

### Taxonomic rank standardization

Retrievable taxonomies were at different ranks, with only 250 (20%) of taxon IDs in the filtered dataset resolvable to species (Fig. S1). Analysing sequence IDs across different ranks confounds ecological patterns with genetic data availability. This is especially true where the specificity of sequence records is taxonomically biased, as our data were here (Fig. S1). However, standardizing taxonomic IDs to a given rank results in either fidelity loss scaling-up species identities, or excluding observations only resolvable to higher ranks. Our main-text analyses used records scaled to genus, allowing us to retain a fine taxonomic grain, with family-level variants included as supplements to ensure results were not driven by excluded data (Figures S1-3). Genus data for analysis comprised 5,870 occurrences of 369 taxa for a total of 11,848,145 sequence reads. These data represent 42.81% of taxa and 58.7% of sequence reads from the filtered dataset (Table S4). Family-level data comprised 8,197 occurrences of 467 taxa for a total of 13,218,264 sequence reads: 54.2% of taxa and 65.5% of sequence reads from the filtered dataset (Table S4).

### Data analysis

#### Richness estimation and model

To assess changes in community composition across ROB age range, we assessed taxon-specific changes in richness as a proxy for community assembly through time. We estimated richness for each taxon group and sample using genus-level presence-absence data, resulting in 486 richness estimates (nine taxon groups × nine ROBs × six pseudoreplicate samples). Richness estimates were used as the response variable in a Poisson-distributed (log-link function) hierarchical Bayesian model (implemented in the brms package; Bürkner, 2017). Models were run with four chains for 10,000 iterations and specified weakly informative priors: normal priors (mu = 0, sigma = 5) for beta coefficients and half-normal priors (mu = 0, sigma = 2) for random effect coefficients. Model assumptions were validated using the DHARMa package (Hartig and Hartig, 2017).

We fitted two hierarchical models to evaluate patterns of taxon-specific richness and their change across the ROB age range. The first was an intercept-only model that contained taxon group as a fixed effect and ROB identity as a random intercept to account for pseudoreplication among samples. The second model added a second fixed effect, ROB age, with initial 9-month surveys treated as a zero-point (with units expressed as months since 9-months). We allowed interactions between ROB age and taxon group to estimate group-specific slopes of richness change over time. We also ran a separate richness model for Mollusca, estimated richness independently for Bivalvia and Gastropoda. All model specification and priors followed those described for the full model above.

#### Shifts in community composition

To assess shifts in community composition across the ROB age range, we complemented the richness analysis with a distance-based redundancy analysis (dbRDA) that tested whether community composition differed based on ROB age. As with the richness models, taxon groups were modelled separately. For each taxon group, genus occurrence within subsampled ROB sequences was converted into a proportional presence matrix. In this matrix, each row was a ROB, each column was a genus, and cells values reflected the proportion of subsamples (out of six) in which a given genus was detected. This is a measure of taxa occurrence over a presumed eDNA sampling filter, with the assumption that a genus detected in all six replicates for a particular ROB was likely in higher abundance than a taxon detected in only one replicate. This provided us with pseudo-abundance data without attempting to translate sequence reads into a proxy for true in-water abundance.

We used each proportional presence matrix as the response variable in a distance-based redundancy analysis (dbrda function, vegan package) (Dixon, 2003), with ROB age as a fixed effect. This analysis constrains compositional variation along fixed effects, with spillover collected in unconstrained dimensions (as per a non-metric multidimensional scaling). The proportion of variation captured by the constrained axes (“constrained inertia”) reflects, in our case, the age-structure of genus composition in the sampled ROBs. Higher constrained inertia reflects greater age-structuring, and vice versa. The significance of age-structuring was determined by comparing observed constrained inertia with null models built from randomly permuted compositional matrices (anova.cca function, vegan package). Taxon groups with high constrained inertia were visualised, with group-specific subgroup classifications added. Again, molluscs were divided into Bivalvia and Gastropoda. Crustacea were split into benthic and non-benthic life history. Flatworms were split into free-living vs parasitic. Fish were divided into small (< 30 cm maximum length) and large (≥30 cm maximum length). Life history and length data were obtained from Global Marine Species Trait Database (Assis and Fragkopoulou, 2024), with genus values determined by the most common category among species of a genus (for categorical traits) and mean maximum length (for fish size).

## Results

### Richness over time

Genus richness varied strongly among taxon groups, describing how the number of taxa present changed across the ROB age range. Differences among taxon groups accounted for most observed variation in richness, with ROB age explaining a smaller additional component. In the taxon-group intercept-only model, fixed effects explained 82.7% of richness variation (marginal R^2^ = 0.827). Including ROB age increased the proportion of variation explained by fixed effects to 85.9% (marginal R^2^ = 0.859), indicating that richness differences among taxa dominated overall patterns, with comparatively smaller age-related effects. Variation attributable to differences among ROBs were minor, with conditional R^2^ (variance explained by both random effects and fixed effects) of 0.871 and 0.906 in the intercept-only and age-slope models respectively.

Molluscs were the richest taxon group, with an estimated twenty genera per sample in a nine-month-old ROB (Fig. 2). Cnidarians and crustaceans had, on average, ten genera in each sample, fish and sponges had seven and segmented worms had five. The remaining three taxon groups, flatworms, roundworms and macroalgae, each had an expected richness of fewer than two genera. For flatworms and roundworms, low estimated richness was likely a function of poor genetic data (Fig. 2, Fig. S1), as fewer than 50% of observations were resolvable to genus. In contrast, all macroalgal sequences were resolved to genus, suggesting poor richness was a result of ecological presence in the sample ROBs or upstream methodological choices (e.g., primer choice) rather than taxonomic uncertainty.

**Figure 2:**
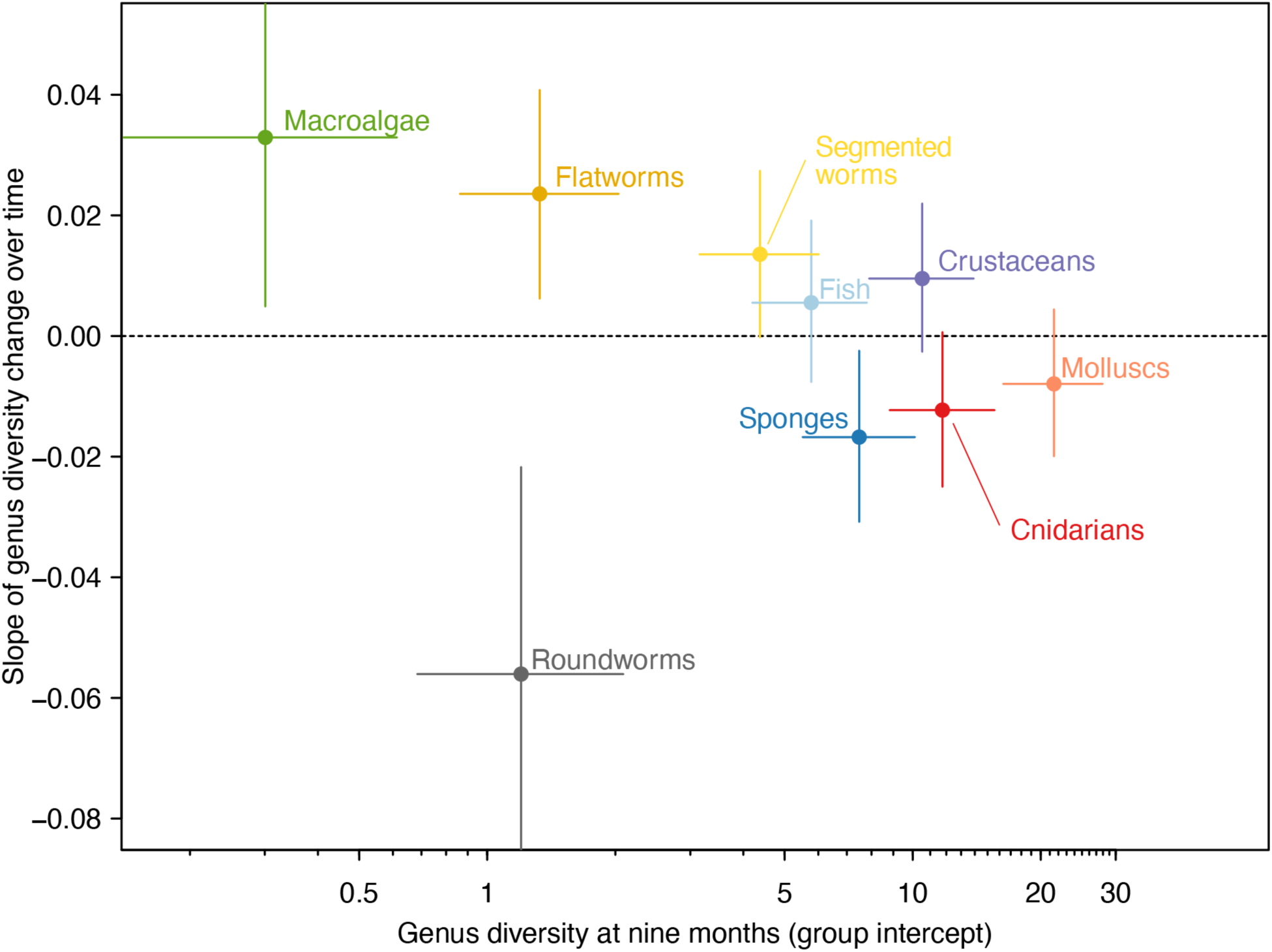
Model coefficients of richness over time model for nine major taxon groups. X-axis position is the estimated genus richness in the earliest age class (nine months-old). Slope of richness change over time is shown on the y-axis. Error bars are 95% confidence intervals.

Five taxon groups exhibited significant changes in richness across the ROB age range. Richness declined with ROB age for cnidarians, sponges and roundworms, whereas flatworms and macroalgae showed increasing richness over time (Fig. 2). With the exception of cnidarians, these temporal trends were consistent when richness was calculated at the family level rather than genus (Fig. S2). Within Mollusca, richness trajectories differed between subgroups. Gastropoda richness decreased in older ROBs, while Bivalvia richness was higher overall and showed little change with ROB age (Fig. 3).

**Figure 3:**
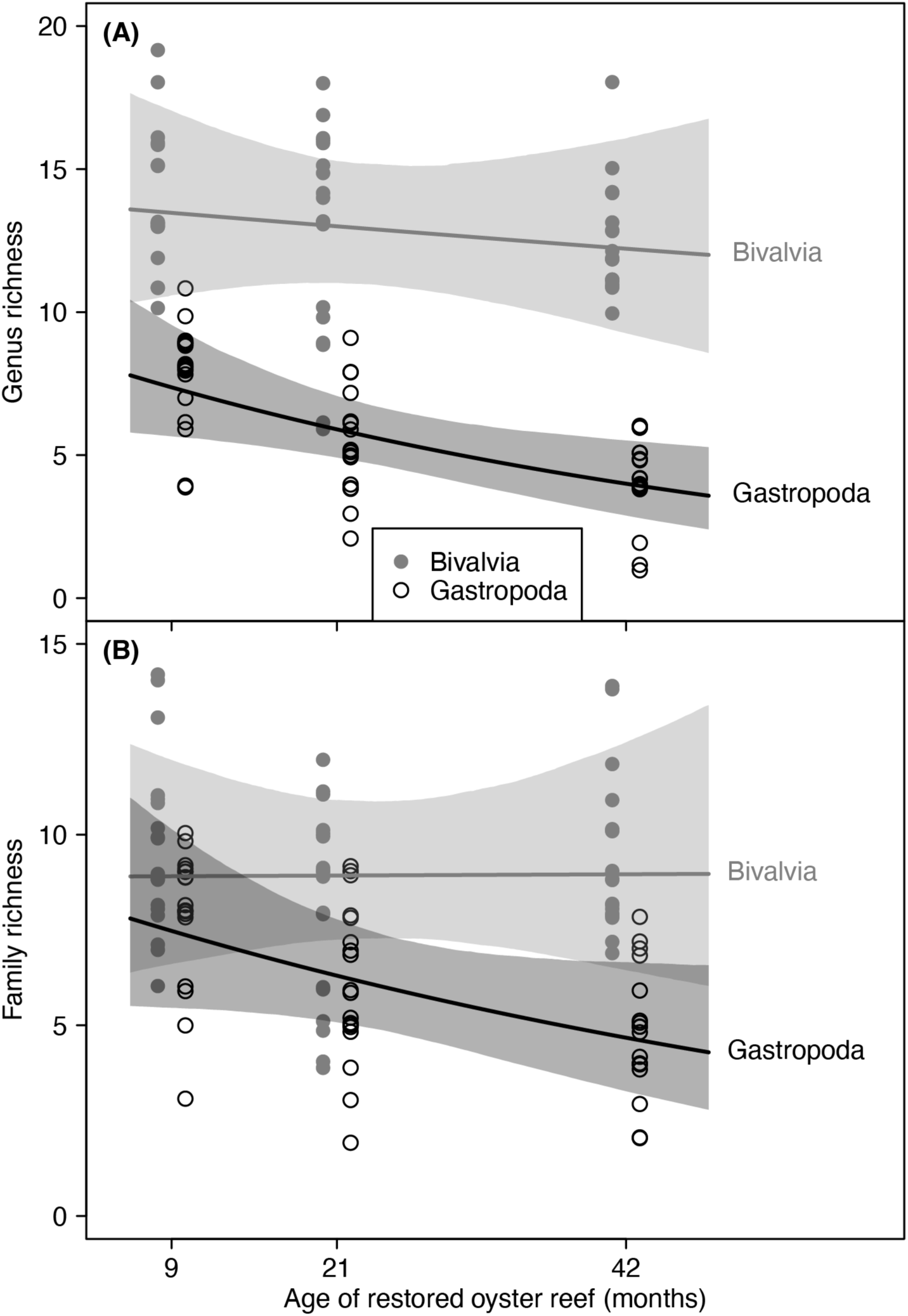
(A) Genus and (B) family richness of two Mollusca classes, Bivalvia and Gastropoda, as a function of age of restored oyster reef. Points are raw richness counts for each sample in each ROB (n = 54 for each class). Lines are mean richness estimates with 95% confidence intervals as shaded regions.

### Compositional shifts

Genus composition was significantly structured by ROB age in six of the nine taxon groups. Flatworm, macroalgae and roundworm groups showed no detectable age-related structuring, consistent with their low genus richness. In taxon groups exhibiting age structuring, similar compositional patterns emerged. Nine-month and 21-month ROBs occupied similar positions on the RDA1 axis, which captured age-related variation in genus composition (Fig. 4). Variation among these younger ROB classes spilled into the first unconstrained MDS axis (y-axis in Fig. 4). Crustaceans, fish, and molluscs also exhibited patterns consistent with convergent succession. Intra-age compositional variation was markedly lower in the 42-month ROBs compared to younger samples (Fig. 4). In the 42-month ROBs, we also observed a trend towards non-benthic crustaceans, larger fish genera and free-living flatworms compared to younger ROBs (Fig. 4).

**Figure 4:**
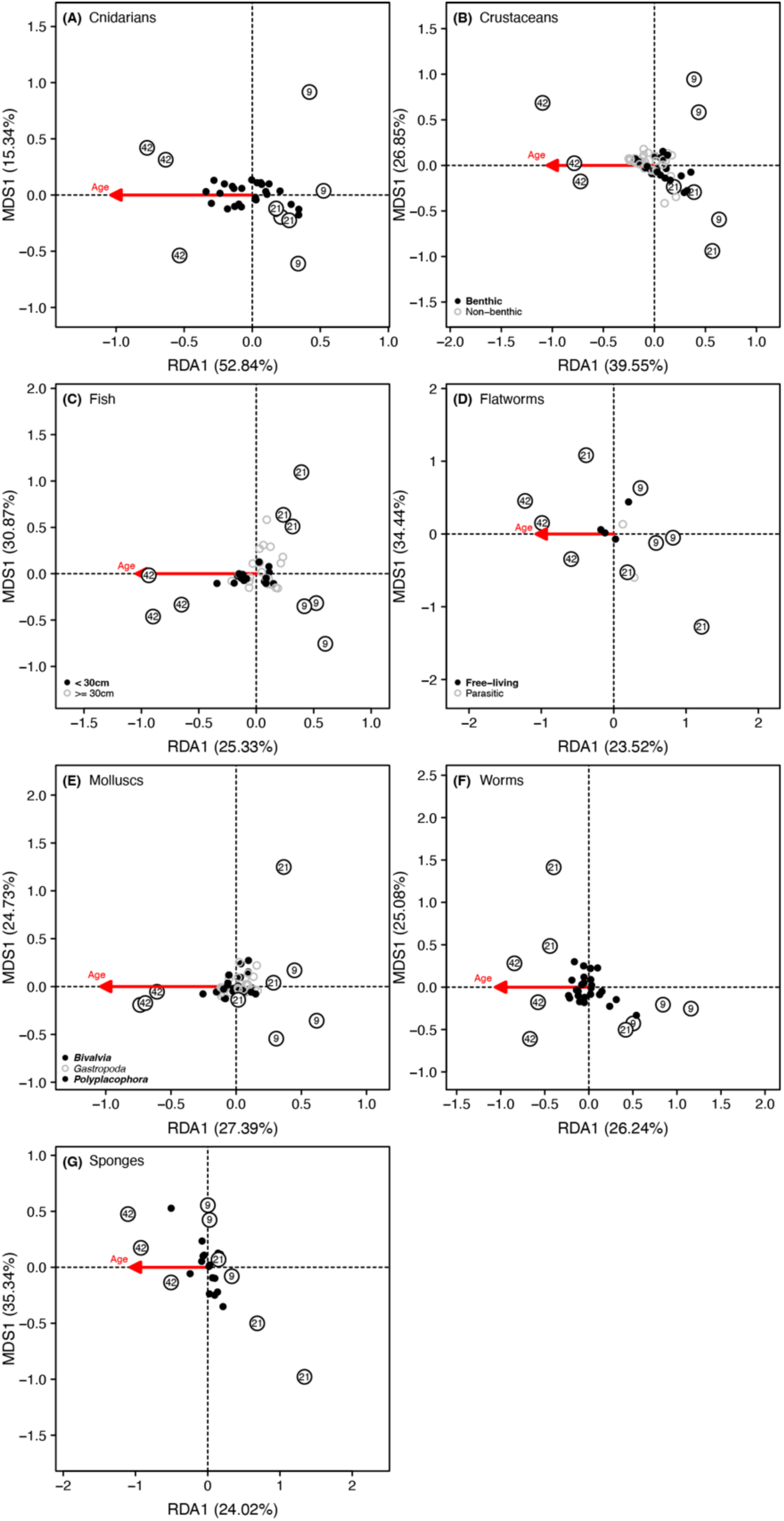
Distance-based redundancy analysis of five taxon groups exhibiting the strongest age-structured composition (Table 1). X-axis is the constrained axis describing compositional variation as a function of ROB age (percentage listed in x-axis label). ROBs are indicated as open circles labelled with age (in months). The direction of the age-structuring is represented by the red arrow. Y-axis is an unconstrained axis that captures the dominant axis of compositional variation after age-structuring was considered. Some taxon groups have additional genus information indicated by point shape.

**Table 1:** Summary of distance-based redundancy analyses of genus composition over time, split by broad taxon group. Constrained inertia reflects compositional variation explainable by ROB age (in months), both in principal coordinate axes and as a proportion of all compositional variation. F-statistic and *p*-value were estimated via comparison with randomly permuted compositional matrices acting as a null hypothesis. Each taxon group was analysed separately.

| Taxon group | Constrained inertia | Constrained inertia (proportion) | pseudo-F | *p*-value |
| --- | --- | --- | --- | --- |
| Cnidarians | 0.240 | 0.542 | 8.278 | 0.001 |
| Crustaceans | 0.501 | 0.414 | 4.954 | 0.001 |
| Fish | 0.224 | 0.270 | 2.589 | 0.022 |
| Flatworms | 0.530 | 0.260 | 2.454 | 0.064 |
| Macroalgae | 0.254 | 0.148 | 1.040 | 0.422 |
| Molluscs | 0.155 | 0.281 | 2.737 | 0.001 |
| Roundworms | 0.336 | 0.328 | 2.439 | 0.133 |
| Segmented worms | 0.328 | 0.247 | 2.298 | 0.009 |
| Sponges | 0.137 | 0.279 | 2.712 | 0.030 |

### Mollusca compositional change

Bivalve molluscs were divided into three broad categories based on frequency of occurrence. Five genera (*Irus*, *Lopha*, *Mytilus*, *Pinctada* and *Saccostrea*) were present in all samples of all ROBs, suggesting a ubiquitous presence. Reflecting their agnostic presence across ROB age, their weighted centroids were central on the age-structured redundancy analysis axis (Fig. 5). We also observed 15 genera that were observed in fewer than 20% of samples. With the exception of *Planostrea*, these were all non-oyster bivalves (primarily saltwater clams). The remaining genera, detected in 20–85% of samples, exhibited the strongest age-structuring (Fig. 5). More of these genera were present in younger versus older ROBs, with only *Modiolus* and *Chama* biased towards detection in 42-month ROBs.

**Figure 5:**
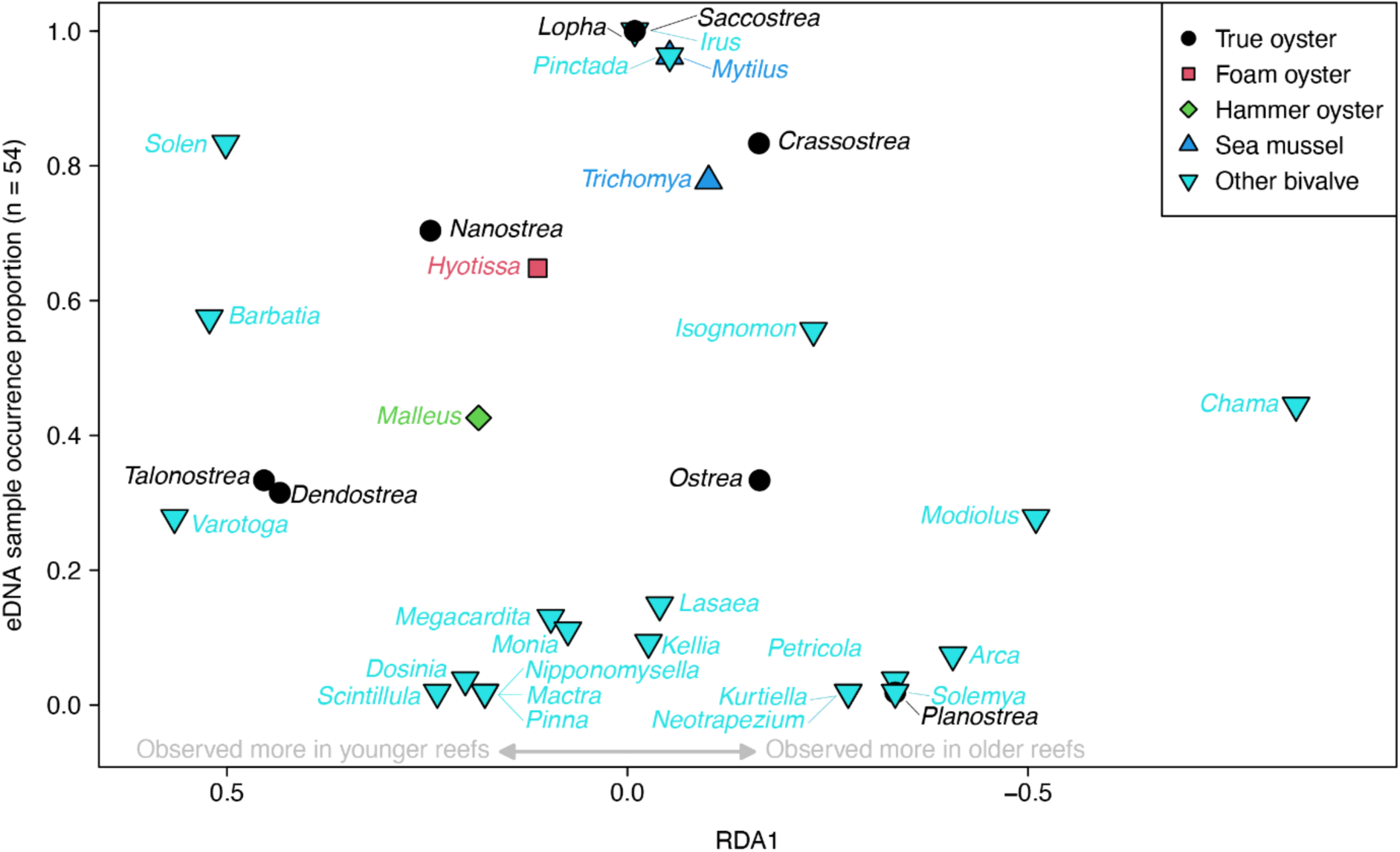
Bivalve genera positions in Mollusca distance-based redundancy analysis (Fig. 4E) as a function of genus presence in eDNA samples (y-axis, proportion out of 54). Genus shape and colour reflect broader bivalve categories, separating oyster and mussel taxa from other bivalves such as clams. Note that x-axis is inverted to read as younger-skewed to older-skewed, following time’s arrow.

## Discussion

### Summary of results

Biodiversity is a common objective of restoration success, with diversity linked to resilience and provision of ecosystem services. We found that modular shellfish restoration structures gained diversity quickly, with genus-level richness in dominant taxonomic groups, particularly molluscs, similar in six-month old structures compared to those retrieved after 48 months. Core reef-forming oyster genera, such as *Saccostrea*, were present in all 54 eDNA samples, evidence that oysters rapidly colonise available hard substrate. This pattern extended beyond foundational taxa to higher trophic habitat users, with crustaceans and fish exhibiting high and consistent richness even in six-month-old reefs, in line with findings from other estuarine habitats (Pierson and Eggleston, 2014). This is a positive sign that, over time, these reefs are recovering to functional ecosystems.

Diversity alone is not a reliable indicator of successful restoration, as richness estimates cannot detect shifts in community composition that may reflect divergent successional trajectories, or the presence of invasive or undesirable taxa. To address this limitation, restoration projects are commonly benchmarked against a known reference state. In the Quandamooka (Moreton Bay) region, however, few shellfish reefs exist following the near total loss of oyster reefs, including the functional extinction of subtidal reef habitats (Diggles, 2013; Gillies et al., 2018). Remnant communities have been used as reference states for biodiversity assessments, but the scale of functional loss of oyster reef ecosystems across continental Australia amid ongoing and intensifying anthropogenic impacts raises uncertainty about whether these remnant reefs truly represent a ‘pre-impact’ state (Cook et al., 2022; Crawford et al., 2020; Gillies et al., 2017). Given this, we relied on emergent ecological patterns to identify potential value-adding or value-removing processes. Our redundancy analyses, which decomposed compositional trajectories based on reef age, provide valuable information to both evaluate restoration success and fingerprint underlying ecological processes. Using this approach, we found that almost all major taxon groups exhibited an age-structured composition, with young reefs more similar to each other than to old reefs, and vice versa.

Compositional patterns identified in this study are evidence of successional processes, with pioneer species being replaced through time. Initial priority effects led to varied compositions in younger reefs, but older reefs exhibited strong compositional similarity. This pattern was consistent for both bivalve molluscs, the dominant habitat-forming organisms in oyster reefs, but also cnidarians, crustaceans and fish, potential evidence of bottom-up trophic structuring. An investigation of remnant oyster reefs in similar environmental conditions could elucidate whether what was found in the 48-months old ROBs samples is effectively a climax community. This represents the best-case scenario for the “field of dream” hypothesis, with newly available substrate rapidly colonized by a diverse assemblage of sessile benthic taxa, with consistent and convergent trajectories across trophic levels. In other regions, active restoration is needed to drive consistent composition in mature oyster reefs (Fariñas-Franco et al., 2013), pointing to the importance of the regional species pool and dispersal connectivity for success in passive restoration efforts. Passive restoration trades-off cost and effort for control; sometimes this has a valuable ecological pay-off, as in our results here, but the rapidity in which dominant taxa colonised our reefs indicates a potential vulnerability to early invasion.

### Limitation of eDNA data

In this study, we demonstrate how eDNA datasets can be used as a fast and inexpensive tool to infer biodiversity at a specific timepoint, and to determine trajectories of biodiversity recovery. However, eDNA has certain limitations which must be accounted for during analysis and interpretation of the data, especially in the context of ecology and conservation (see e.g. Beng and Corlett, 2020 and Gold et al., 2021 for a more detailed discussion). Diversity estimates and ecological compositions constructed from eDNA data must be interpreted cautiously. Observed diversities are a mix of the diversity of organisms living in, on, and near the structure at the time, spatial and temporal bleeding of broader taxa presence in nearby habitats or organisms that were present at an earlier time. We believe that sampling modular reef structures in tanks, rather than the surrounding water column is likely to preferentially recover eDNA from taxa that were repeatedly present on or interation with the reef. While DNA originating from outside reef residents may be concentrated by filter feeders; evidence suggests that the fate of this eDNA may depend on filtration and retention pathways. For example, (Jeunen et al., 2023) showed that vertebrate eDNA was inconsistently and weakly recovered from bivalve gill tissue, in contrast to robust detection from water and sponge samples from the same environment.

In contrast, a preprint by (Bolte et al., 2025) demonstrated that the Pacific oyster (*Magallana gigas)* can retain target organism DNA within digestive tissues following exposure, suggesting that eDNA retention in bivalves may depend on the tissue sampled and the mechanism of uptake (ingestion vs passive filtration). Interesting, Friebertshauser et al., 2019 showed that bivalve filtration can significantly reduce eDNA concentrations in the surrounding water column in a density-dependent manner, indicating that bivalves may effectively remove eDNA from the environment even when that DNA is not subsequently recoverable from gill tissues.

Comparing species diversity detected from samples within hours or ROBS arrival in the laboratory with those taken after a defined depuration period, may provide a means of distinguishing which taxa are genuinely associated with ROBS versus those that are transient, loosely attached, or derived from adjacent habitats. This temporal comparison could help distinguish resident communities from opportunistic or incidental species and improve interpretation of eDNA signals derived from ROBS-based sampling.

### Matching sequences with reference databases

Diversity estimates from eDNA data are constrained by incompleteness and structure of the available reference databases. Only taxa represented in the databases can be detected. In this study, resolvable taxonomic rank of eDNA sequences varied substantially among taxon groups, likely reflecting broader “Linnean shortfalls”, where many taxa have not been formally described (Hortal et al., 2015). Just over half of all sequences in our dataset were not resolvable to any taxonomic rank, despite similar nucleotide length to sequences with valid taxonomy (irresolvable μ = 102.5 vs resolvable μ = 110), indicating that database coverage rather than sequence quality was the primary limitation. Even among resolvable sequences, taxonomic resolution was highly group specific. We know far more about fish than we do worms, complicating attempts to detect diversity trends across both groups simultaneously. Although published guidelines discuss how to reliably curate eDNA data to assign taxonomy (Alberdi et al., 2018), they offer limited guidelines to handle biased or uneven assignments when estimating ecological metrics like diversity. Treating molecular OTUs as direct surrogates for ecological diversity, or scaling sequences to a standard rank, both have fundamental issues. The first does not provide any ecological information outside of DNA diversity, a “richness of everything”, and conflates genome size and primer fidelity with diversity. Combining taxonomic assignments, such as class and species, treats ranks with different volumes of ecological and evolutionary information as equivalent units of diversity. Even filtering data into a single rank, like we did here, is not a complete solution. Our diversity results are incomparable across taxa groups; it is impossible to know whether the mollusc assemblage in our reefs was more diverse than the sponge assemblage, or whether it was a function of upstream biases in genetic databases. We treated eDNA data cautiously, maximizing data availability while avoiding cross-taxa comparisons, using convergent within-group compositional patterns used as potential evidence of ecological processes.

These challenges are further compounded by variability in reference database construction: while custom databases can be assembled from open-access repositories such as GenBank and Bold (Benson et al., 2013; Ratnasingham and Hebert, 2007), and curating the data. The use of these online repositories to create a personalised database can lead to multimodal distributions of sequence similarity that can be due to a variety of factors (e.g. hybridization between congeners, misidentification of congeners) (Locatelli et al., 2020). Therefore, level of species assignment improves with locally derived curated databases (Blackman et al., 2023), specific to the area in which the study is conducted, and with a higher sequence similarity. Ongoing initiatives by authors and others to expand the genomic reference library for Moreton Bay/Quandamooka should progressively enhance the resolution and reliability of species-level assignments for eDNA samples collected in this region.

### Estimations of species abundance

One of the major limitations of the specific assay used in this research paper, is the inability to estimate species abundance and biomass of the various taxa. Indeed, the dataset can be only interpreted as presence/absence (occurrence). While occurrence datasets are a valuable tool, particularly in the context of detection of threatened and invasive species (e.g. in Moreton Bay for White’s seahorse, *Hippocampus whitei* and Suminoe oyster, *Magallana ariakensis*, respectively), and have also been used to monitor populations at large spatial scales and to identify high value habitats, abundance data could give a better insight of the structure of the community and the status of the population (Beng and Corlett, 2020; Vörös et al., 2017; Weltz et al., 2017).

### Contamination risk

Lastly, sample contamination risk is a generally high in eDNA studies and can occur at various stages of sample handling (Sepulveda et al., 2020). To minimize contamination, the tubs which were used to collect the ROBs were sterilized with the use of bleach, but as their dimensions were quite large, it’s likely that the extensive area captured DNA particles present in the air. Moreover, the tubs were brought along on the boat to the ROBs’ deployment site on a boat, and to and from boat to the ROBs site by walking in intertidal zone. We tried to minimize the tubs exposure to external contamination by always keeping the lids on, but during the ROBs retrieval, after sterilization. Our approach accounted for these risks by trying to replicate the same experimental procedures for both the samples and the negative controls, which allowed us to manage potential false positives through statistical methods.

## Conclusions

This study demonstrates that eDNA can be used as a tool to infer biodiversity gains and track ecological recovery states in modular shellreef restoration projects. By applying a taxonomic rank standardization approach, we were able to analyse complex metabarcoding datasets and detect both rapid increases in richness and more gradual, directional changes in community composition through time. These patterns indicate that eDNA is capable of capturing key aspects of community assembly and successional development on restored reefs.

However, while eDNA provides a powerful tool for quantifying ecological trajectories, assessing restoration success or failure may ultimately depend on the availability of appropriate reference states or clearly defined targets. In the absence of such benchmarks, eDNA-based assessments are best suited to tracking relative change and convergence through time rather than determining absolute recovery. Nevertheless, our results highlight the potential of eDNA as a scalable and non-invasive approach for monitoring restoration outcomes, supporting its use in evaluating the effectiveness of reef-building interventions such as modular shellfish reefs.

## Data availability statement

Data and R code necessary to reproduce results are available at 10.5281/zenodo.18690589.

## References

Alberdi, A., Aizpurua, O., Gilbert, M.T.P., Bohmann, K., 2018. Scrutinizing key steps for reliable metabarcoding of environmental samples. Methods Ecol. Evol. 9, 134–147. 10.1111/2041-210X.12849

Aronson, J., Alexander, S., 2013. Steering Towards Sustainability Requires More Ecological Restoration. Nat. Conserv. 11, 127–137. DOI%2010.4322/natcon.2013.021

Assis, J., Fragkopoulou, E., 2024. marineSpeciesTraits: Global Marine Species Traits Database. 10.6084/m9.figshare.27620676.v3

Baggett, L.P., Powers, S.P., Brumbaugh, R.D., Coen, L.D., DeAngelis, B.M., Greene, J.K., Hancock, B.T., Morlock, S.M., Allen, B.L., Breitburg, D.L., Bushek, D., Grabowski, J.H., Grizzle, R.E., Grosholz, E.D., La Peyre, M.K., Luckenbach, M.W., McGraw, K.A., Piehler, M.F., Westby, S.R., zu Ermgassen, P.S.E., 2015. Guidelines for evaluating performance of oyster habitat restoration. Restor. Ecol. 23, 737–745. 10.1111/rec.12262

Balensiefer, M., Rossi, R., Ardinghi, N., Cenni, M., Ugolini, M., 2004. SER international primer on ecological restoration. Soc. Ecol. Restor. Wash.

Beck, M.W., Brumbaugh, R.D., Airoldi, L., Carranza, A., Coen, L.D., Crawford, C., Defeo, O., Edgar, G.J., Hancock, B., Kay, M.C., Lenihan, H.S., Luckenbach, M.W., Toropova, C.L., Zhang, G.F., Guo, X.M., 2011. Oyster Reefs at Risk and Recommendations for Conservation, Restoration, and Management. Bioscience 61, 107–116. 10.1525/bio.2011.61.2.5

Benayas, J.M.R., Newton, A.C., Diaz, A., Bullock, J.M., 2009. Enhancement of Biodiversity and Ecosystem Services by Ecological Restoration: A Meta-Analysis. Science 325, 1121–1124. 10.1126/science.1172460

Beng, K.C., Corlett, R.T., 2020. Applications of environmental DNA (eDNA) in ecology and conservation: opportunities, challenges and prospects. Biodivers. Conserv. 29, 2089–2121. 10.1007/s10531-020-01980-0

Benson, D.A., Cavanaugh, M., Clark, K., Karsch-Mizrachi, I., Lipman, D.J., Ostell, J., Sayers, E.W., 2013. GenBank. Nucleic Acids Res. 41, D36–D42. 10.1093/nar/gks1195

Blackman, R.C., Walser, J.-C., Rüber, L., Brantschen, J., Villalba, S., Brodersen, J., Seehausen, O., Altermatt, F., 2023. General principles for assignments of communities from eDNA: Open versus closed taxonomic databases. Environ. DNA 5, 326–342. 10.1002/edn3.382

Bolte, B., Bissett, A., Craw, P., McDougall, C., Wynne, J.W., 2025. Use of the Pacific oyster (Magallana gigas) as a natural sampler for the detection of Neoparamoeba perurans. 10.1101/2025.08.06.669027

Brewer, J.S., Menzel, T., 2009. A Method for Evaluating Outcomes of Restoration When No Reference Sites Exist. Restor. Ecol. 17, 4–11. 10.1111/j.1526-100X.2008.00456.x

Bürkner, P.-C., 2017. brms: An R Package for Bayesian Multilevel Models Using Stan. J. Stat. Softw. 80, 1–28. 10.18637/jss.v080.i01

Chamberlain, S., Vanhoorne, B., 2023. worrms: World Register of Marine Species (WoRMS) Client_. R package version 0.4.3,.

Cole, J.R., Wang, Q., Fish, J.A., Chai, B., McGarrell, D.M., Sun, Y., Brown, C.T., Porras-Alfaro, A., Kuske, C.R., Tiedje, J.M., 2014. Ribosomal Database Project: data and tools for high throughput rRNA analysis. Nucleic Acids Res. 42, D633–D642. 10.1093/nar/gkt1244

Cook, P.A., Warnock, B., Gillies, C.L., Hams, A.B., 2022. Historical abundance and distribution of the native flat oyster (Ostrea angasi) in estuaries of the Great Southern region of Western Australia help to prioritise potential sites for contemporary oyster reef restoration. Mar. Freshw. Res. 73, 48–56. 10.1071/Mf21058

Crawford, C., Edgar, G., Gillies, C.L., Heller-Wagner, G., 2020. Relationship of biological communities to habitat structure on the largest remnant flat oyster reef (Ostrea angasi) in Australia. Mar. Freshw. Res. 71, 972–983. 10.1071/Mf19171

Diggles, B., 2013. Historical epidemiology indicates water quality decline drives loss of oyster (Saccostrea glomerata) reefs in Moreton Bay, Australia. N. Z. J. Mar. Freshw. Res. 47, 561–581. 10.1080/00288330.2013.781511

Dixon, P., 2003. VEGAN, a package of R functions for community ecology. J. Veg. Sci. 14, 927–930. 10.1111/j.1654-1103.2003.tb02228.x

Duarte, C.M., Conley, D.J., Carstensen, J., Sánchez-Camacho, M., 2009. Return to Neverland: Shifting Baselines Affect Eutrophication Restoration Targets. Estuaries Coasts 32, 29–36. 10.1007/s12237-008-9111-2

Edgar, R.C., 2016. SINTAX: a simple non-Bayesian taxonomy classifier for 16S and ITS sequences. biorxiv 074161.

Eger, A.M., Earp, H.S., Friedman, K., Gatt, Y., Hagger, V., Hancock, B., Kaewsrikhaw, R., Mcleod, E., Moore, A.M., Niner, H.J., Razafinaivo, F., Sousa, A., Stankovic, M., Worthington, T.A., Bayraktarov, E., Saunders, M., Vergés, A., Reeves, S., 2022. The need, opportunities, and challenges for creating a standardized framework for marine restoration monitoring and reporting. Biol. Conserv. 266. ARTN%20109429%2010.1016/j.biocon.2021.109429

Fariñas-Franco, J.M., Allcock, L., Smyth, D., Roberts, D., 2013. Community convergence and recruitment of keystone species as performance indicators of artificial reefs. J. Sea Res. 78, 59–74. 10.1016/j.seares.2012.10.008

Friebertshauser, R., Shollenberger, K., Janosik, A., Garner, J.T., Johnston, C., 2019. The effect of bivalve filtration on eDNA-based detection of aquatic organisms. PLOS ONE 14, e0222830. 10.1371/journal.pone.0222830

Fukami, T., Nakajima, M., 2011. Community assembly: alternative stable states or alternative transient states? Ecol. Lett. 14, 973–984. 10.1111/j.1461-0248.2011.01663.x

Gibbs, M., Ross, P., Scanes, E., Gibbs, J., Rotolo-Ross, R., Parker, L., 2023. Extending conservation of coastal and oyster reef restoration for First Nations cultural revitalization. Conserv Biol.

Gibbs, M.C., Parker, L.M., Scanes, E., Ross, P.M., 2024. Recognising the importance of shellfish to First Nations peoples, Indigenous and Traditional Ecological Knowledge in aquaculture and coastal management in Australia. Mar. Freshw. Res. 75, NULL-NULL.

Gillies, C.L., Crawford, C., Hancock, B., 2017. Restoring Angasi oyster reefs: What is the endpoint ecosystem we are aiming for and how do we get there? Ecol. Manag. Restor. 18, 214–222. 10.1111/emr.12278

Gillies, C.L., McLeod, I.M., Alleway, H.K., Cook, P., Crawford, C., Creighton, C., Diggles, B., Ford, J., Hamer, P., Heller-Wagner, G., Lebrault, E., Le Port, A., Russell, K., Sheaves, M., Warnock, B., 2018. Australian shellfish ecosystems: Past distribution, current status and future direction. Plos One 13. ARTN%20e0190914%2010.1371/journal.pone.0190914

Gold, Z., Sprague, J., Kushner, D.J., Marin, E.Z., Barber, P.H., 2021. eDNA metabarcoding as a biomonitoring tool for marine protected areas. PLOS ONE 16, e0238557. 10.1371/journal.pone.0238557

Grabowski, J.H., Peterson, C.H., 2007. Restoring Oyster Reefs to Recover Ecosystem Services. Ecosyst. Eng. Plants Protists 281–298.

Hagger, V., Vanderklift, M.A., Beaumont, K.P., Olds, A., Dittman, S., Lovelock, C.E., 2025. Biodiversity gains can be achieved with coastal wetland restoration for plants, invertebrates, birds, and bats. Ecol. Indic. 178, 113805. 10.1016/j.ecolind.2025.113805

Hallett, L.M., Diver, S., Eitzel, M.V., Olson, J.J., Ramage, B.S., Sardinas, H., Statman-Weil, Z., Suding, K.N., 2013. Do We Practice What We Preach? Goal Setting for Ecological Restoration. Restor. Ecol. 21, 312–319. 10.1111/rec.12007

Hartig, F., Hartig, M.F., 2017. Package ‘dharma.’ R Package 531, 532.

Hilderbrand, R.H., Watts, A.C., Randle, A.M., 2005. The myths of restoration ecology. Ecol. Soc. 10.

Hobbs, R.J., Harris, J.A., 2001. Restoration ecology: Repairing the Earth’s ecosystems in the new millennium. Restor. Ecol. 9, 239–246. DOI%2010.1046/j.1526-100x.2001.009002239.x

Hortal, J., Bello, F. de, Diniz-Filho, J.A.F., Lewinsohn, T.M., Lobo, J.M., Ladle, R.J., 2015. Seven Shortfalls that Beset Large-Scale Knowledge of Biodiversity. Annu. Rev. Ecol. Evol. Syst. 46, 523–549. 10.1146/annurev-ecolsys-112414-054400

Jeunen, G.-J., Cane, J.S., Ferreira, S., Strano, F., von Ammon, U., Cross, H., Day, R., Hesseltine, S., Ellis, K., Urban, L., Pearson, N., Olmedo-Rojas, P., Kardailsky, A., Gemmell, N.J., Lamare, M., 2023. Assessing the utility of marine filter feeders for environmental DNA (eDNA) biodiversity monitoring. Mol. Ecol. Resour. 23, 771–786. 10.1111/1755-0998.13754

Locatelli, N.S., McIntyre, P.B., Therkildsen, N.O., Baetscher, D.S., 2020. GenBank’s reliability is uncertain for biodiversity researchers seeking species-level assignment for eDNA. Proc. Natl. Acad. Sci. 117, 32211–32212. 10.1073/pnas.2007421117

Lotze, H.K., Lenihan, H.S., Bourque, B.J., Bradbury, R.H., Cooke, R.G., Kay, M.C., Kidwell, S.M., Kirby, M.X., Peterson, C.H., Jackson, J.B.C., 2006. Depletion, Degradation, and Recovery Potential of Estuaries and Coastal Seas. Science 312, 1806–1809. 10.1126/science.1128035

Myers, R.A., Worm, B., 2003. Rapid worldwide depletion of predatory fish communities. Nature 423, 280–283. 10.1038/nature01610

Ogburn, D.M., White, I., Mcphee, D.P., 2007. The Disappearance of Oyster Reefs from Eastern Australian Estuaries—Impact of Colonial Settlement or Mudworm Invasion? Coast. Manag. 35, 271–287. 10.1080/08920750601169618

Palmer, M.A., Ambrose, R.F., Poff, N.L., 1997. Ecological theory and community restoration ecology. Restor. Ecol. 5, 291–300. DOI%2010.1046/j.1526-100X.1997.00543.x

Pauly, D., 1995. Anecdotes and the shifting baseline syndrome of fisheries. Trends Ecol. Evol. 10, 430.

Perring, M.P., Standish, R.J., Price, J.N., Craig, M.D., Erickson, T.E., Ruthrof, K.X., Whiteley, A.S., Valentine, L.E., Hobbs, R.J., 2015. Advances in restoration ecology: rising to the challenges of the coming decades. Ecosphere 6. Artn%20131%2010.1890/Es15-00121.1

Pinnegar, J.K., Engelhard, G.H., 2008. The ‘shifting baseline’ phenomenon: a global perspective. Rev. Fish Biol. Fish. 18, 1–16. 10.1007/s11160-007-9058-6

R Core Team, 2021. R: A Language and Environment for Statistical Computing. R Foundation for Statistical Computing, Vienna, Austria.

Ratnasingham, S., Hebert, P.D.N., 2007. bold: The Barcode of Life Data System (http://www.barcodinglife.org). Mol. Ecol. Notes 7, 355–364. 10.1111/j.1471-8286.2007.01678.x

Ruiz-Jaen, M.C., Mitchell Aide, T., 2005. Restoration Success: How Is It Being Measured? Restor. Ecol. 13, 569–577. 10.1111/j.1526-100X.2005.00072.x

Saenz-Agudelo, P., Delrieu-Trottin, E., DiBattista, J.D., Martínez-Rincon, D., Morales-González, S., Pontigo, F., Ramírez, P., Silva, A., Soto, M., Correa, C., 2022. Monitoring vertebrate biodiversity of a protected coastal wetland using eDNA metabarcoding. Environ. DNA 4, 77–92. 10.1002/edn3.200

Sepulveda, A.J., Hutchins, P.R., Forstchen, M., Mckeefry, M.N., Swigris, A.M., 2020. The Elephant in the Lab (and Field): Contamination in Aquatic Environmental DNA Studies. Front. Ecol. Evol. 8. 10.3389/fevo.2020.609973

Smith, R.S., Lusk, B., Castorani, M.C.N., 2022. Restored oyster reefs match multiple functions of natural reefs within a decade. Conserv. Lett. 15. ARTN%20e12883%2010.1111/conl.12883

Suding, K.N., 2011. Toward an Era of Restoration in Ecology: Successes, Failures, and Opportunities Ahead. Annu. Rev. Ecol. Evol. Syst. Vol 42 42, 465–487. 10.1146/annurev-ecolsys-102710-145115

Suding, K.N., Gross, K.L., Houseman, G.R., 2004. Alternative states and positive feedbacks in restoration ecology. Trends Ecol. Evol. 19, 46–53. 10.1016/j.tree.2003.10.005

Vörös, J., Márton, O., Schmidt, B.R., Gál, J.T., Jelić, D., 2017. Surveying Europe’s Only Cave-Dwelling Chordate Species (Proteus anguinus) Using Environmental DNA. PLOS ONE 12, e0170945. 10.1371/journal.pone.0170945

Walker, L.R., Wardle, D.A., Bardgett, R.D., Clarkson, B.D., 2010. The use of chronosequences in studies of ecological succession and soil development. J. Ecol. 98, 725–736. 10.1111/j.1365-2745.2010.01664.x

Weltz, K., Lyle, J.M., Ovenden, J., Morgan, J.A.T., Moreno, D.A., Semmens, J.M., 2017. Application of environmental DNA to detect an endangered marine skate species in the wild. PLOS ONE 12, e0178124. 10.1371/journal.pone.0178124

Wilkinson, S., 2025. Laboratory methods for Wilderlab ECOSCAN panels (Technical note). Wilderlab NZ Ltd., Wellington, New Zealand.

Wilkinson, S.P., Davy, S.K., Bunce, M., Stat, M., 2018. Taxonomic identification of environmental DNA with informatic sequence classification trees. PeerJ Prepr.

Wilkinson, S.P., Gault, A.A., Welsh, S.A., Smith, J.P., David, B.O., Hicks, A.S., Fake, D.R., Suren, A.M., Shaffer, M.R., Jarman, S.N., Bunce, M., 2024. TICI: a taxon-independent community index for eDNA-based ecological health assessment. PeerJ 12, e16963. 10.7717/peerj.16963

zu Ermgassen, P.S.E., Spalding, M.D., Grizzle, R.E., Brumbaugh, R.D., 2013. Quantifying the Loss of a Marine Ecosystem Service: Filtration by the Eastern Oyster in US Estuaries. Estuaries Coasts 36, 36–43. 10.1007/s12237-012-9559-y

