## Supplementary Data for "From deployment to functioning ecosystems: assessing taxonomic richness and community composition on modular shellfish reefs using environmental DNA metabarcoding"

**Table S1** Sample information for the first round of sampling on 08/05/2024.

| **Sample #** | **Sample ID** | **Collection Date** | **Latitude** | **Longitude** | **Volume Filtered (mL)** | **Age of deployment** | **Assays** |
| --- | --- | --- | --- | --- | --- | --- | --- |
| 543762 | ROBCONTROL 1A | 2024-05-08 | NA | NA | 900 | NA | WV,RV,LG,XI,HD,BE,BU,UM,WG,TP,MZ,GV,BX,GD,WO,RJ |
| 543766 | ROBCONTROL 1B | 2024-05-08 | NA | NA | 1000 | NA | WV,RV,LG,XI,HD,BE,BU,UM,WG,TP,MZ,GV,BX,GD,WO,RJ |
| 543743 | ROBCONTROL 1C | 2024-05-08 | NA | NA | 850 | NA | WV,RV,LG,XI,HD,BE,BU,UM,WG,TP,MZ,GV,BX,GD,WO,RJ |
| 543759 | ROB400-1-1 | 2024-05-08 | -27.458940 | 153.397200 | 650 | 48 months | WV,RV,LG,XI,HD,BE,BU,UM,WG,TP,MZ,GV,BX,GD,WO,RJ |
| 543758 | ROB400-1-2 | 2024-05-08 | -27.458940 | 153.397200 | 700 | 48 months | WV,RV,LG,XI,HD,BE,BU,UM,WG,TP,MZ,GV,BX,GD,WO,RJ |
| 543756 | ROB400-1-3 | 2024-05-08 | -27.458940 | 153.397200 | 1000 | 48 months | WV,RV,LG,XI,HD,BE,BU,UM,WG,TP,MZ,GV,BX,GD,WO,RJ |
| 543755 | ROB400-1-4 | 2024-05-08 | -27.458940 | 153.397200 | 850 | 48 months | WV,RV,LG,XI,HD,BE,BU,UM,WG,TP,MZ,GV,BX,GD,WO,RJ |
| 543750 | ROB400-1-5 | 2024-05-08 | -27.458940 | 153.397200 | 1000 | 48 months | WV,RV,LG,XI,HD,BE,BU,UM,WG,TP,MZ,GV,BX,GD,WO,RJ |
| 543761 | ROB400-1-6 | 2024-05-08 | -27.458940 | 153.397200 | 800 | 48 months | WV,RV,LG,XI,HD,BE,BU,UM,WG,TP,MZ,GV,BX,GD,WO,RJ |
| 543764 | ROB400-2-1 | 2024-05-08 | -27.458940 | 153.397200 | 600 | 48 months | WV,RV,LG,XI,HD,BE,BU,UM,WG,TP,MZ,GV,BX,GD,WO,RJ |
| 543760 | ROB400-2-2 | 2024-05-08 | -27.458940 | 153.397200 | 600 | 48 months | WV,RV,LG,XI,HD,BE,BU,UM,WG,TP,MZ,GV,BX,GD,WO,RJ |
| 543763 | ROB400-2-3 | 2024-05-08 | -27.458940 | 153.397200 | 750 | 48 months | WV,RV,LG,XI,HD,BE,BU,UM,WG,TP,MZ,GV,BX,GD,WO,RJ |
| 543768 | ROB400-2-4 | 2024-05-08 | -27.458940 | 153.397200 | 900 | 48 months | WV,RV,LG,XI,HD,BE,BU,UM,WG,TP,MZ,GV,BX,GD,WO,RJ |
| 543767 | ROB400-2-5 | 2024-05-08 | -27.458940 | 153.397200 | 1000 | 48 months | WV,RV,LG,XI,HD,BE,BU,UM,WG,TP,MZ,GV,BX,GD,WO,RJ |
| 543757 | ROB400-2-6 | 2024-05-08 | -27.458940 | 153.397200 | 1000 | 48 months | WV,RV,LG,XI,HD,BE,BU,UM,WG,TP,MZ,GV,BX,GD,WO,RJ |
| 543751 | ROB400-3-1 | 2024-05-08 | -27.458940 | 153.397200 | 1000 | 48 months | WV,RV,LG,XI,HD,BE,BU,UM,WG,TP,MZ,GV,BX,GD,WO,RJ |
| 543749 | ROB400-3-2 | 2024-05-08 | -27.458940 | 153.397200 | 1000 | 48 months | WV,RV,LG,XI,HD,BE,BU,UM,WG,TP,MZ,GV,BX,GD,WO,RJ |
| 543752 | ROB400-3-3 | 2024-05-08 | -27.458940 | 153.397200 | 1000 | 48 months | WV,RV,LG,XI,HD,BE,BU,UM,WG,TP,MZ,GV,BX,GD,WO,RJ |
| 543754 | ROB400-3-4 | 2024-05-08 | -27.458940 | 153.397200 | 850 | 48 months | WV,RV,LG,XI,HD,BE,BU,UM,WG,TP,MZ,GV,BX,GD,WO,RJ |
| 543747 | ROB400-3-5 | 2024-05-08 | -27.458940 | 153.397200 | 820 | 48 months | WV,RV,LG,XI,HD,BE,BU,UM,WG,TP,MZ,GV,BX,GD,WO,RJ |
| 543742 | ROB400-3-6 | 2024-05-08 | -27.458940 | 153.397200 | 900 | 48 months | WV,RV,LG,XI,HD,BE,BU,UM,WG,TP,MZ,GV,BX,GD,WO,RJ |

**Table S2** Sample information for the second round of sampling on 29/08/2024.

| **Sample #** | **Sample ID** | **Collection Date** | **Latitude** | **Longitude** | **Volume Filtered (mL)** | **Age of deployment** | **Assays** |
| --- | --- | --- | --- | --- | --- | --- | --- |
| 546747 | ROB4-1 | 2024-08-29 | -27.4589400 | 153.3972000 | 1000 | 6 month | WV,RV,LG,XI,HD,BE,BU,UM,WG,TP,MZ,GV,BX,GD,WO,RJ |
| 546760 | ROB4-2 | 2024-08-29 | -27.4589400 | 153.3972000 | 1000 | 6 month | WV,RV,LG,XI,HD,BE,BU,UM,WG,TP,MZ,GV,BX,GD,WO,RJ |
| 546732 | ROB4-3 | 2024-08-29 | -27.4589400 | 153.3972000 | 1000 | 6 month | WV,RV,LG,XI,HD,BE,BU,UM,WG,TP,MZ,GV,BX,GD,WO,RJ |
| 546746 | ROB4-4 | 2024-08-29 | -27.4589400 | 153.3972000 | 1000 | 6 month | WV,RV,LG,XI,HD,BE,BU,UM,WG,TP,MZ,GV,BX,GD,WO,RJ |
| 546778 | ROB4-5 | 2024-08-29 | -27.4589400 | 153.3972000 | 1000 | 6 month | WV,RV,LG,XI,HD,BE,BU,UM,WG,TP,MZ,GV,BX,GD,WO,RJ |
| 546758 | ROB4-6 | 2024-08-29 | -27.4589400 | 153.3972000 | 1000 | 6 month | WV,RV,LG,XI,HD,BE,BU,UM,WG,TP,MZ,GV,BX,GD,WO,RJ |
| 546767 | ROB5-1 | 2024-08-29 | -27.4589400 | 153.3972000 | 850 | 18 months | WV,RV,LG,XI,HD,BE,BU,UM,WG,TP,MZ,GV,BX,GD,WO,RJ |
| 546785 | ROB5-2 | 2024-08-29 | -27.4589400 | 153.3972000 | 1000 | 18 months | WV,RV,LG,XI,HD,BE,BU,UM,WG,TP,MZ,GV,BX,GD,WO,RJ |
| 546759 | ROB5-3 | 2024-08-29 | -27.4589400 | 153.3972000 | 1000 | 18 months | WV,RV,LG,XI,HD,BE,BU,UM,WG,TP,MZ,GV,BX,GD,WO,RJ |
| 546788 | ROB5-4 | 2024-08-29 | -27.4589400 | 153.3972000 | 1000 | 18 months | WV,RV,LG,XI,HD,BE,BU,UM,WG,TP,MZ,GV,BX,GD,WO,RJ |
| 546769 | ROB5-5 | 2024-08-29 | -27.4589400 | 153.3972000 | 900 | 18 months | WV,RV,LG,XI,HD,BE,BU,UM,WG,TP,MZ,GV,BX,GD,WO,RJ |
| 546768 | ROB5-6 | 2024-08-29 | -27.4589400 | 153.3972000 | 900 | 18 months | WV,RV,LG,XI,HD,BE,BU,UM,WG,TP,MZ,GV,BX,GD,WO,RJ |
| 546798 | ROB6-1 | 2024-08-29 | -27.4589400 | 153.3972000 | 900 | 6 month | WV,RV,LG,XI,HD,BE,BU,UM,WG,TP,MZ,GV,BX,GD,WO,RJ |
| 546757 | ROB6-2 | 2024-08-29 | -27.4589400 | 153.3972000 | 1000 | 6 month | WV,RV,LG,XI,HD,BE,BU,UM,WG,TP,MZ,GV,BX,GD,WO,RJ |
| 546774 | ROB6-3 | 2024-08-29 | -27.4589400 | 153.3972000 | 1000 | 6 month | WV,RV,LG,XI,HD,BE,BU,UM,WG,TP,MZ,GV,BX,GD,WO,RJ |
| 546761 | ROB6-4 | 2024-08-29 | -27.4589400 | 153.3972000 | 700 | 6 month | WV,RV,LG,XI,HD,BE,BU,UM,WG,TP,MZ,GV,BX,GD,WO,RJ |
| 546787 | ROB6-5 | 2024-08-29 | -27.4589400 | 153.3972000 | 1000 | 6 month | WV,RV,LG,XI,HD,BE,BU,UM,WG,TP,MZ,GV,BX,GD,WO,RJ |
| 546765 | ROB6-6 | 2024-08-29 | -27.4589400 | 153.3972000 | 900 | 6 month | WV,RV,LG,XI,HD,BE,BU,UM,WG,TP,MZ,GV,BX,GD,WO,RJ |
| 546764 | ROB7-1 | 2024-08-29 | -27.4589400 | 153.3972000 | 850 | 18 months | WV,RV,LG,XI,HD,BE,BU,UM,WG,TP,MZ,GV,BX,GD,WO,RJ |
| 546784 | ROB7-2 | 2024-08-29 | -27.4589400 | 153.3972000 | 950 | 18 months | WV,RV,LG,XI,HD,BE,BU,UM,WG,TP,MZ,GV,BX,GD,WO,RJ |
| 546781 | ROB7-3 | 2024-08-29 | -27.4589400 | 153.3972000 | 950 | 18 months | WV,RV,LG,XI,HD,BE,BU,UM,WG,TP,MZ,GV,BX,GD,WO,RJ |
| 546771 | ROB7-4 | 2024-08-29 | -27.4589400 | 153.3972000 | 950 | 18 months | WV,RV,LG,XI,HD,BE,BU,UM,WG,TP,MZ,GV,BX,GD,WO,RJ |
| 546792 | ROB7-5 | 2024-08-29 | -27.4589400 | 153.3972000 | 950 | 18 months | WV,RV,LG,XI,HD,BE,BU,UM,WG,TP,MZ,GV,BX,GD,WO,RJ |
| 546766 | ROB7-6 | 2024-08-29 | -27.4589400 | 153.3972000 | 950 | 18 months | WV,RV,LG,XI,HD,BE,BU,UM,WG,TP,MZ,GV,BX,GD,WO,RJ |
| 546776 | ROB8-1 | 2024-08-29 | -27.4589400 | 153.3972000 | 900 | 18 months | WV,RV,LG,XI,HD,BE,BU,UM,WG,TP,MZ,GV,BX,GD,WO,RJ |
| 546790 | ROB8-2 | 2024-08-29 | -27.4589400 | 153.3972000 | 950 | 18 months | WV,RV,LG,XI,HD,BE,BU,UM,WG,TP,MZ,GV,BX,GD,WO,RJ |
| 546791 | ROB8-3 | 2024-08-29 | -27.4589400 | 153.3972000 | 950 | 18 months | WV,RV,LG,XI,HD,BE,BU,UM,WG,TP,MZ,GV,BX,GD,WO,RJ |
| 546772 | ROB8-4 | 2024-08-29 | -27.4589400 | 153.3972000 | 950 | 18 months | WV,RV,LG,XI,HD,BE,BU,UM,WG,TP,MZ,GV,BX,GD,WO,RJ |
| 546794 | ROB8-5 | 2024-08-29 | -27.4589400 | 153.3972000 | 950 | 18 months | WV,RV,LG,XI,HD,BE,BU,UM,WG,TP,MZ,GV,BX,GD,WO,RJ |
| 546797 | ROB8-6 | 2024-08-29 | -27.4589400 | 153.3972000 | 950 | 18 months | WV,RV,LG,XI,HD,BE,BU,UM,WG,TP,MZ,GV,BX,GD,WO,RJ |
| 546752 | ROB9-1 | 2024-08-29 | -27.4589400 | 153.3972000 | 850 | 6 month | WV,RV,LG,XI,HD,BE,BU,UM,WG,TP,MZ,GV,BX,GD,WO,RJ |
| 546777 | ROB9-2 | 2024-08-29 | -27.4589400 | 153.3972000 | 560 | 6 month | WV,RV,LG,XI,HD,BE,BU,UM,WG,TP,MZ,GV,BX,GD,WO,RJ |
| 546742 | ROB9-3 | 2024-08-29 | -27.4589400 | 153.3972000 | 600 | 6 month | WV,RV,LG,XI,HD,BE,BU,UM,WG,TP,MZ,GV,BX,GD,WO,RJ |
| 546786 | ROB9-4 | 2024-08-29 | -27.4589400 | 153.3972000 | 1000 | 6 month | WV,RV,LG,XI,HD,BE,BU,UM,WG,TP,MZ,GV,BX,GD,WO,RJ |
| 546779 | ROB9-5 | 2024-08-29 | -27.4589400 | 153.3972000 | 1000 | 6 month | WV,RV,LG,XI,HD,BE,BU,UM,WG,TP,MZ,GV,BX,GD,WO,RJ |
| 546751 | ROB9-6 | 2024-08-29 | -27.4589400 | 153.3972000 | 1000 | 6 month | WV,RV,LG,XI,HD,BE,BU,UM,WG,TP,MZ,GV,BX,GD,WO,RJ |
| 546762 | Control1 | 2024-08-29 | NA | NA | 800 | NA | WV,RV,LG,XI,HD,BE,BU,UM,WG,TP,MZ,GV,BX,GD,WO,RJ |
| 546795 | Control2 | 2024-08-29 | NA | NA | 800 | NA | WV,RV,LG,XI,HD,BE,BU,UM,WG,TP,MZ,GV,BX,GD,WO,RJ |
| 546796 | Control3 | 2024-08-29 | NA | NA | 800 | NA | WV,RV,LG,XI,HD,BE,BU,UM,WG,TP,MZ,GV,BX,GD,WO,RJ |

**Table S3** List of metabarcoding primers used to target a wide range of taxonomic groups in Wilderlab NZ Ltd. assays for environmental DNA investigations (*indicates primers were developed in-house at Wilderlab.

| Assay | Primer | Sequence | Annealing (°C) | Length | Notes | Reference | Original Name | Link |
| --- | --- | --- | --- | --- | --- | --- | --- | --- |
| **BE** | BEF | CCCTGCCHTTTGTACACAC | 52 | 19 | General eukaryotes 18S V9 | Amaral-Zettler et al. 2009 | 1380F | 10.1371/journal.pone.0006372 |
|  | BUR | CCTTCYGCAGGTTCACCTAC | 52 | 20 | General eukaryotes 18S V9 | Amaral-Zettler et al. 2009 | 1510R | 10.1371/journal.pone.0006372 |
| **BU** | BUF | TTGTACACACCGCCC | 52 | 15 | General eukaryotes 18S V9 | Amaral-Zettler et al. 2009 | 1389F | 10.1371/journal.pone.0006372 |
|  | BUR | CCTTCYGCAGGTTCACCTAC | 52 | 20 | General eukaryotes 18S V9 | Amaral-Zettler et al. 2009 | 1510R | 10.1371/journal.pone.0006372 |
| **BX** | BXF | GCCAGTAGTCATATGCTTGTCT | 52 | 22 | General eukaryote 18S | Pochon et al. 2013 | 18S_uni_1F | https://doi.org/10.1371/journal.pone.0073935 |
|  | BXR | GCCTGCTGCCTTCCTT | 52 | 16 | General eukaryote 18S | Pochon et al. 2013 | 18S_uni_400R | https://doi.org/10.1371/journal.pone.0073935 |
| **GD** | GDF | GARTCTTTGAACGCAAATGGC | 52 | 21 | Coral ITS2 | Brian et al. 2019 | SCLER28SRev | https://link.springer.com/article/10.1007/s00338-018-01762-9 |
|  | GCR | GCTTATTAATATGCTTAAATTCAGCG | 52 | 26 | Coral ITS2 | Alexander et al. 2019 | Acro874_Rev | https://link.springer.com/article/10.1007/s00338-018-01762-9 |
| **GV** | GVF | GTGAACCTGCGGAAGGATC | 52 | 19 | Vascular plants ITS1 | Andree et al. 2011 | MicroSSU | doi:10.1128/AEM.01978-10 |
|  | GVR | GATATCCGTTGCCGAGAGTC | 52 | 20 | Vascular plants ITS1 | Baamrane et al. 2012 | ITS1-F | https://doi.org/10.1371/journal.pone.0035643 |
| **HD** | HDF | GGACGATAAGACCCTATAAA | 52 | 20 | Crustaceans 16S | Komai et al. 2019 | MiDeca-F | https://mbmg.pensoft.net/article/33835/ |
|  | HDR | ACGCTGTTATCCCTAAAGT | 52 | 19 | Crustaceans 16S | Komai et al. 2019 | MiDeca-R | <https://mbmg.pensoft.net/article/33835/> |
| **LG** | LGF* | CGGCGTAAAGWGTGGTTAGG | 58 | 20 | Fish mt12S | In house |  |  |
|  | LGR | CATAGTGGGGTATCTAATCCCAGTTTG | 58 | 16 | Fish mt12S | Miya et al. 2015 | MiFish-U-R | https://doi.org/10.1098/rsos.150088 |
| **MZ** | MZF | CTTCTTCAGGTGGAACTCCAG | 52 | 21 | Vascular plants rbcL | Bradley et al. 2007 | rbcL19b | DOI 10.1002/ajp.20384 |
|  | MZR | GTCACCACAAACAGAGACTAAAGCAAGT | 52 | 28 | Vascular plants rbcL | Modif. from Bradley et al. 2007 | rbcLZ1 (modified) | DOI 10.1002/ajp.20384 |
| **RV** | RVF | TTAGATACCCCACTATGC | 58 | 18 | Vertebrate 12S ecoprimers | Riaz et al. 2011 | 12S-V5-F | https://doi.org/10.1093/nar/gkr732 |
|  | RVR | TAGAACAGGCTCCTCTAG | 58 | 18 | Vertebrate 12S ecoprimers | Riaz et al. 2011 | 12S-V5-R | https://doi.org/10.1093/nar/gkr732 |
| **TP** | TPF | GGGCAATCCTGAGCCAA | 52 | 17 | Vascular plants trnL | Taberlet et al. 2006 | trnL-g | https://doi.org/10.1093/nar/gkl938 |
|  | TPR | CCATTGAGTCTCTGCACCTATC | 52 | 22 | Vascular plants trnL | Taberlet et al. 2006 | trnL-h | https://doi.org/10.1093/nar/gkl938 |
| **UM** | UMF | GGATTAGATACCCTGGTA | 52 | 18 | Bacteria 16S | Morey et al. 2006 | F785e | https://doi.org/10.1128/jcm.00670-06 |
|  | UMR | CCGTCAATTCMTTTRAGTTT | 52 | 20 | Bacteria 16S | Lane et al. 1985 | 907r | https://doi.org/10.1073/pnas.82.20.6955 |
| **WG** | WGF | TTDTAAAAGMCGAGAAGACCC | 52 | 20 | Venerid clams 16S | Prié et al. 2021 | Vene01_F | https://link.springer.com/article/10.1007/s10750-020-04260-8 |
|  | WGR | CSCTGTTATCCCYRCGGTA | 52 | 19 | Venerid clams 16S | Prié et al. 2021 | Vene01_R | https://link.springer.com/article/10.1007/s10750-020-04260-8 |
| **WO** | WOF | AAGAAGACCCCGTGCARC | 58 | 18 | Oysters 16S | Dugal et al. 2024 | NOyster16SF | https://doi.org/10.1016/j.ecss.2024.108722 |
|  | WOR | CTGTTATCCCCGGCGTAAC | 58 | 19 | Oysters 16S | Dugal et al. 2024 | NOyster16SR | https://doi.org/10.1016/j.ecss.2024.108722 |
| **WV** | WVF | GACGAGAAGACCCTWTGGAGC | 58 | 21 | Vertebrate 16S | Based on Nester et al. 2020 | 16S_FishSyn_Short forward | <https://doi.org/10.1002/edn3.93> |
|  | WVR | CCRYGGTCGCCCCAAC | 58 | 16 | Vertebrate 16S | Based on Nester et al. 2020 | 16S_FishSyn_Short reverse | <https://doi.org/10.1002/edn3.93> |

**Table S4:** Table of sequence read data during each stage of filtering. Observations were the occurrence of a taxa in an eDNA sample. Sequence reads represent individual read counts from upstream bioinformatic processing. Taxa were unique IDs from the amplicon sequence variants cross-referenced with Genbank and RDP (see methods for details).

|  | **Data filtering step** | | | **Observation count** | **Sequence read count** | **Taxa count** |
| --- | --- | --- | --- | --- | --- | --- |
| 1 | Full sequence read data | | | 423,152 | 60,378,284 | 3,698 |
| 2 | Remove "root" entries | | | 204,305 | 47,360,023 | 3,697 |
| 3 | Remove non-marine/brackish taxa using WORMs | | | 182,750 | 41,750,439 | 2,257 |
| 4 | Remove taxa from undesired groups (e.g., bacteria, diatoms) | | | 20,959 | 20,274,494 | 873 |
| 5 | Remove contaminant taxa (using controls) | | | 20,935 | 20,268,475 | 864 |
| 6 | Remove control samples from data | | | 20,743 | 20,196,640 | 862 |
| 7a | Retain reads resolvable to genus | | | 5,870 | 11,848,145 | 369 |
| 7b | Retain reads resolvable to family | | | 8,197 | 13,218,264 | 467 |

**Table S5:** Table of contamination sequence reads in control and sample replicates. This table was used to include or exclude taxa for subsequent analyses, using dataset at filtering step 4 (Table S4). Taxa were excluded if they were present in at least one sample in both control rounds, with the exception of Saccostrea.

| **Taxonomic ID** | **Taxon group** | **Rank** | **Presence in eDNA samples** | | | | **Keep or exclude** |
| --- | --- | --- | --- | --- | --- | --- | --- |
|  |  |  | **Round 1 controls** | **Round 2 controls** | **Round 1 samples** | **Round 2 samples** |  |
| Hexanauplia | Crustaceans | class | 3 | 1 | 18 | 36 | exclude |
| *Saccostrea* | Molluscs | genus | 3 | 1 | 18 | 36 | keep |
| Actiniaria | Cnidarians | order | 3 | 0 | 18 | 36 | keep |
| Anthoathecata | Cnidarians | order | 3 | 0 | 18 | 34 | keep |
| Anthozoa | Cnidarians | class | 3 | 0 | 18 | 36 | keep |
| *Bougainvillia* | Cnidarians | genus | 3 | 0 | 7 | 4 | keep |
| Bougainvilliidae | Cnidarians | family | 3 | 0 | 6 | 3 | keep |
| Calanoida | Crustaceans | order | 3 | 0 | 15 | 33 | keep |
| Cephalaspidea | Molluscs | order | 3 | 0 | 5 | 5 | keep |
| *Euterpina* | Crustaceans | genus | 3 | 0 | 15 | 13 | keep |
| Harpacticoida | Crustaceans | order | 3 | 0 | 18 | 32 | keep |
| Hydrozoa | Cnidarians | class | 3 | 0 | 18 | 36 | keep |
| *Microstomum* | Flatworms | genus | 3 | 0 | 4 | 18 | keep |
| *Notomastus* | Worms | genus | 3 | 0 | 18 | 36 | keep |
| *Obelia bidentata* | Cnidarians | species | 3 | 0 | 17 | 36 | keep |
| Ostreidae | Molluscs | family | 3 | 0 | 18 | 36 | keep |
| Phyllodocida | Worms | order | 1 | 2 | 13 | 33 | exclude |
| *Sertularia* | Cnidarians | genus | 3 | 0 | 18 | 36 | keep |
| Arcoidea | Molluscs | superfamily | 2 | 0 | 15 | 16 | keep |
| Campanularia | Cnidarians | genus | 2 | 0 | 2 | 9 | keep |
| Chromadorea | Roundworms | class | 2 | 0 | 15 | 28 | keep |
| Gastropoda | Molluscs | class | 2 | 0 | 18 | 36 | keep |
| Harpacticidae | Crustaceans | family | 2 | 0 | 1 | 13 | keep |
| *Mictyris longicarpus* | Crustaceans | species | 2 | 0 | 2 | 0 | keep |
| Podocopida | Crustaceans | order | 2 | 0 | 17 | 36 | keep |
| Polychaeta | Worms | class | 2 | 0 | 18 | 35 | keep |
| Rhabditophora | Flatworms | class | 2 | 0 | 3 | 11 | keep |
| Rhabdocoela | Flatworms | order | 2 | 0 | 8 | 15 | keep |
| *Acanthopagrus australis* | Fish | species | 1 | 0 | 18 | 17 | keep |
| *Acartia* | Crustaceans | genus | 1 | 0 | 16 | 11 | keep |
| *Aetobatus narinari* | Fish | species | 1 | 0 | 0 | 2 | keep |
| *Alepes kleinii* | Fish | species | 1 | 0 | 0 | 0 | keep |
| *Bestiolina* | Crustaceans | genus | 1 | 0 | 3 | 12 | keep |
| Calcarea | Sponges | class | 1 | 0 | 17 | 36 | keep |
| *Caloglossa* | Macroalgae | genus | 1 | 0 | 1 | 3 | keep |
| Capitellidae | Worms | family | 1 | 0 | 18 | 23 | keep |
| Cirratulidae | Worms | family | 1 | 0 | 2 | 31 | keep |
| Cyclopidae | Crustaceans | family | 1 | 0 | 0 | 0 | keep |
| Cytheruridae | Crustaceans | family | 1 | 0 | 0 | 0 | keep |
| Dactylogyridea | Flatworms | order | 1 | 0 | 3 | 0 | keep |
| Dactylopusiidae | Crustaceans | family | 1 | 0 | 15 | 30 | keep |
| Dugesiidae | Flatworms | family | 1 | 0 | 0 | 0 | keep |
| *Eusyllis* | Worms | genus | 1 | 0 | 0 | 0 | keep |
| *Gerres* | Fish | genus | 1 | 0 | 11 | 8 | keep |
| *Gymnothorax pseudothyrsoideus* | Fish | species | 1 | 0 | 1 | 0 | keep |
| *Halicephalobus* | Roundworms | genus | 1 | 0 | 1 | 0 | keep |
| *Haliclona* | Sponges | genus | 1 | 0 | 9 | 32 | keep |
| *Hymeniacidon* | Sponges | genus | 1 | 0 | 9 | 29 | keep |
| *Hyperietta* | Crustaceans | genus | 1 | 0 | 3 | 0 | keep |
| *Irus crenatus* | Molluscs | species | 0 | 1 | 18 | 36 | keep |
| Leptothecata | Cnidarians | order | 1 | 0 | 18 | 36 | keep |
| *Lopha* | Molluscs | genus | 1 | 0 | 18 | 36 | keep |
| *Mycale* | Sponges | genus | 1 | 0 | 9 | 35 | keep |
| *Myrianida* | Worms | genus | 1 | 0 | 11 | 3 | keep |
| Paraonidae | Worms | family | 1 | 0 | 0 | 0 | keep |
| *Pinctada* | Molluscs | genus | 1 | 0 | 18 | 34 | keep |
| *Pionosyllis* | Worms | genus | 1 | 0 | 0 | 0 | keep |
| *Pomatoceros* | Worms | genus | 1 | 0 | 17 | 21 | keep |
| *Rastrelliger kanagurta* | Fish | species | 1 | 0 | 1 | 0 | keep |
| Spionidae | Worms | family | 1 | 0 | 18 | 35 | keep |
| Syllidae | Worms | family | 1 | 0 | 18 | 24 | keep |
| Thecostraca | Crustaceans | class | 1 | 0 | 2 | 10 | keep |
| Thoracica | Crustaceans | infraclass | 1 | 0 | 15 | 27 | keep |
| Tricladida | Flatworms | order | 1 | 0 | 0 | 0 | keep |

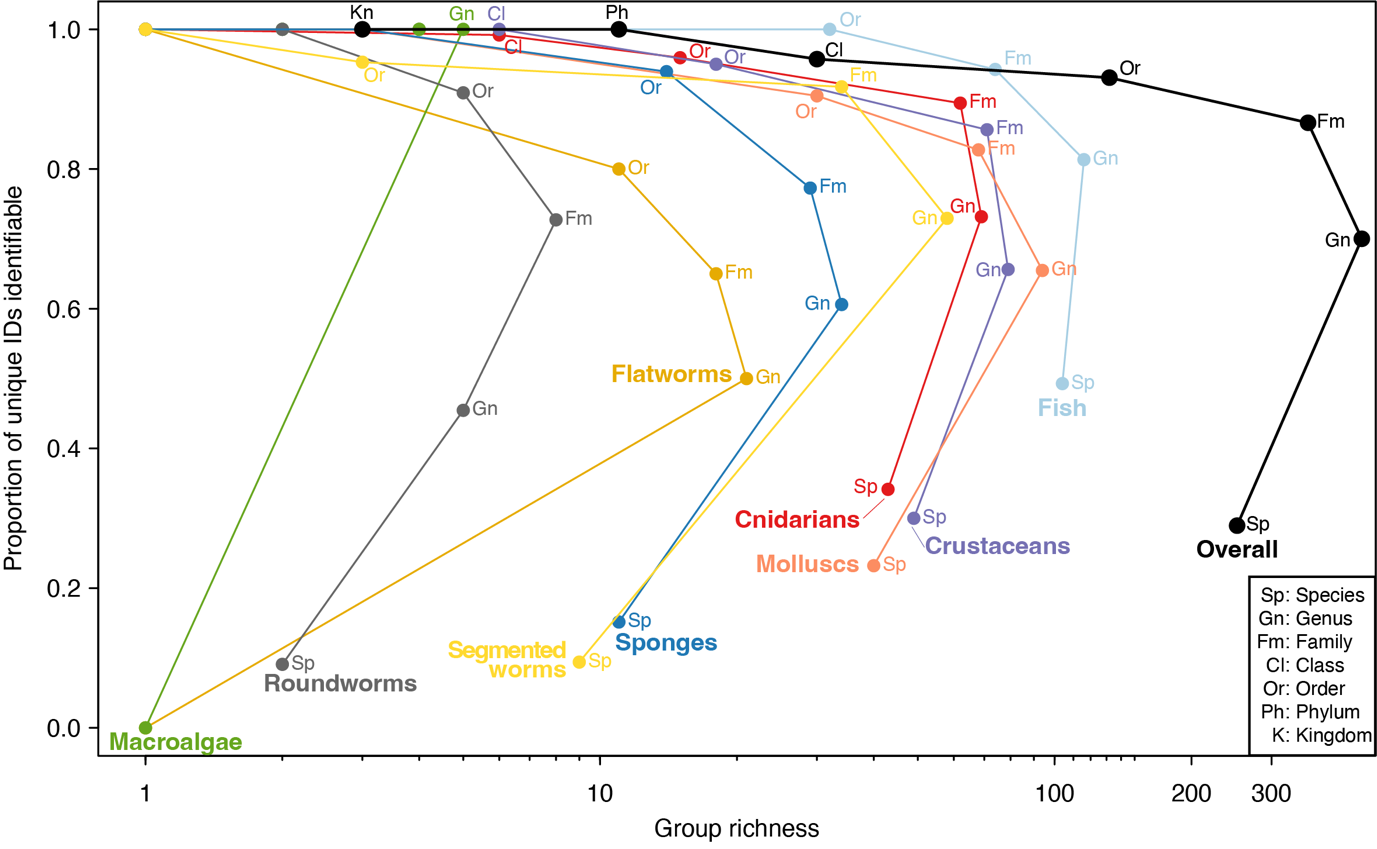

**Figure S1:** Total richness of taxon groups in eDNA data (x-axis) at different taxonomic ranks, with the proportion of unique taxon IDs resolvable to each rank indicated on the y-axis. Overall, eDNA data was richest at the genus rank, with 70% of taxon IDs resolvable to genus. This was strongly biased by taxon, with some groups like fish, with > 80% of IDs resolvable to genus, while roundworms were < 50%. This figure captures the trade-off between taxonomic specificity (via lower ranks) and data inclusion (via proportion of IDs resolvable to a given rank). In addition, the strong division on the x-axis reflects systematic biases in both taxonomic and genetic data, which, if analysed together, would be conflated with true sample richness.

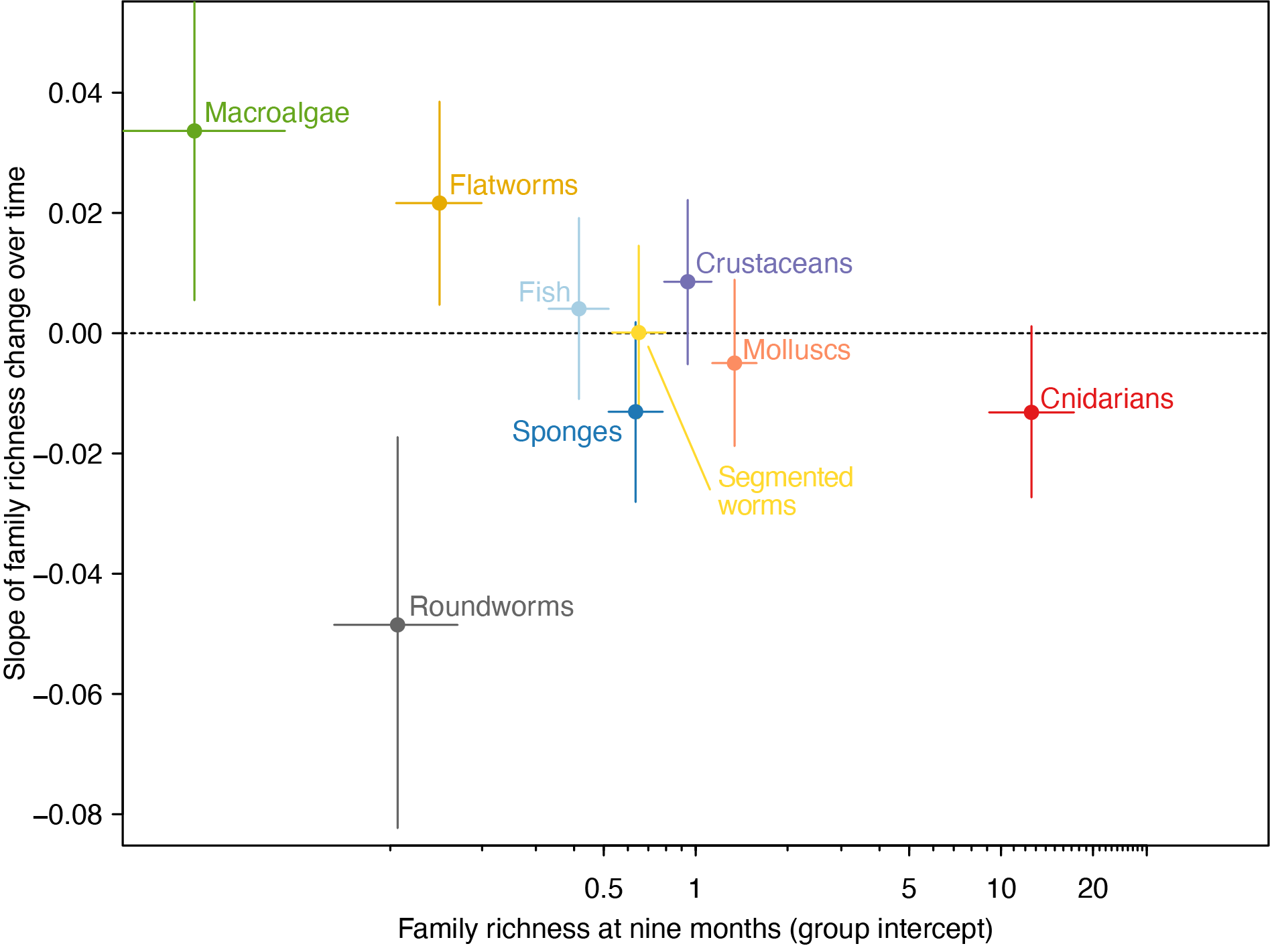

**Figure S2:** **Version of main text Figure 1 analyzed using family-level taxonomic identifications.** Model coefficients of richness over time model for nine major taxon groups. X-axis position is the estimated genus richness in the earliest age class (six months-old). Slope of richness change over time is shown on the y-axis. Error bars are 95% confidence intervals.

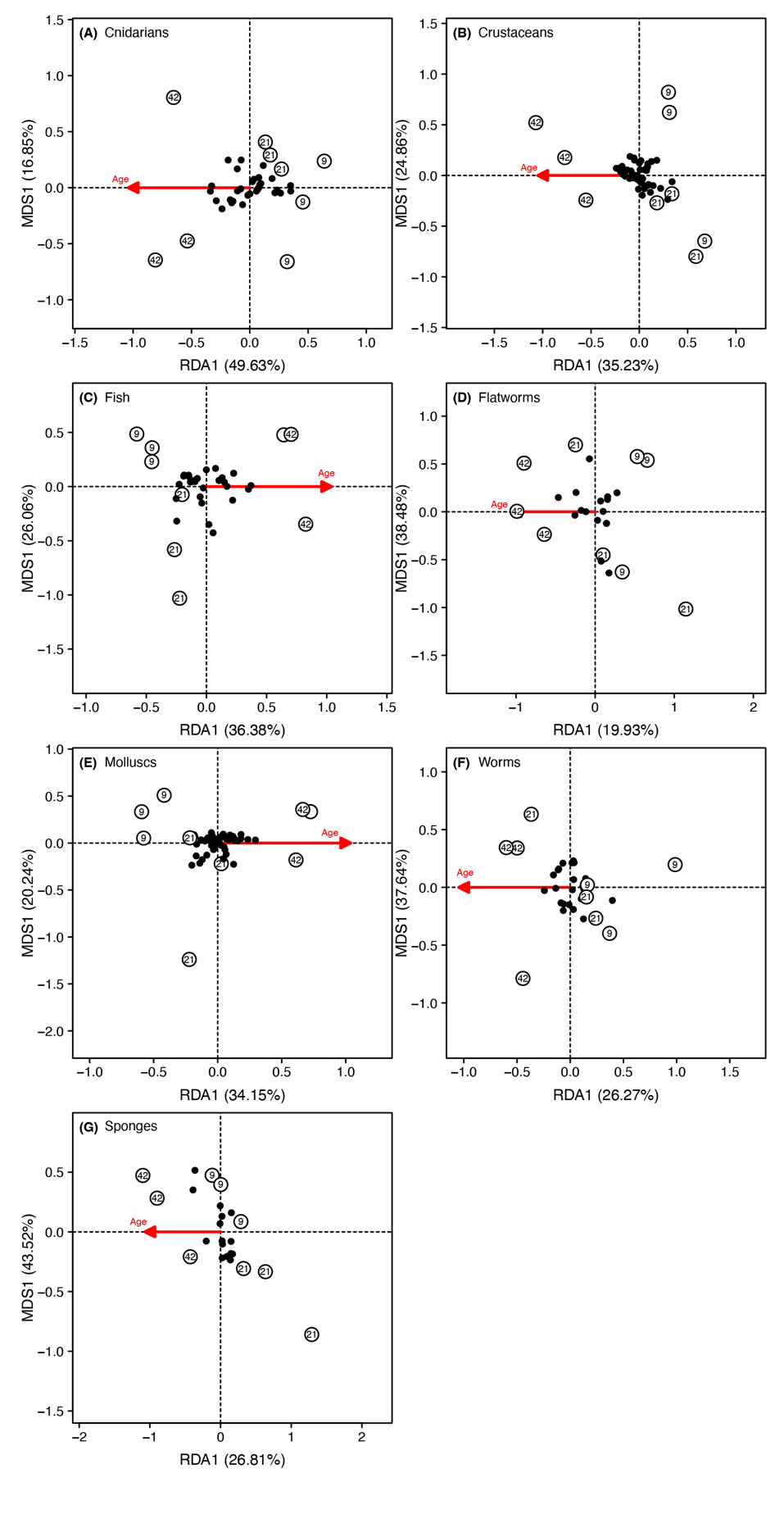

**Figure S3:** **Version of main text Figure 3 analyzed using family-level taxonomic identifications**. Distance-based redundancy analysis of five taxon groups exhibiting the strongest age-structured composition (Table 1). X-axis is the constrained axis describing compositional variation as a function of ROB age (percentage listed in x-axis label). ROBs are indicated as open circles labelled with age (in months). The direction of the age-structuring is represented by the red arrow. Y-axis is an unconstrained axis that captures the dominant axis of compositional variation after age-structuring was considered. Some taxon groups have additional genus information indicated by point shape.

Wilderlab Methods
